# Histone H3K27 methylation states are sequentially catalyzed in cycling cells

**DOI:** 10.64898/2026.04.30.721988

**Authors:** Jacob E. Greene, Kami Ahmad, Steven Henikoff

## Abstract

Polycomb domains form at silenced genes and are marked by histone H3 lysine 27 tri-methylation, but it is unclear how this methylation is maintained following DNA replication in proliferating cells. Here, we use CUT&Tag chromatin profiling to track the dynamics of H3K27 methylation in human cells. We find that tri-methylation in Polycomb domains is maintained by stepwise methylation after DNA replication. Outside of Polycomb domains, thousands of inactive genes gain histone H3K27 di-methylation hours after DNA replication. Acute treatment with small-molecule inhibitors of the Polycomb Repressive Complex 2 (PRC2) histone methyltransferase slows H3K27 methylation during S-phase of the cell cycle and increases acetylation of the residue. This effect is most pronounced at Polycomb domains replicating in early S-phase, suggesting that early replication and slow histone methylation confers plasticity to developmentally silenced genes. Together, our results explain how H3K27 modification patterns are faithfully duplicated in rapidly proliferating cells.

## Introduction

In different cell contexts genes are transcriptionally active or inactive, and these states of gene activity must be stably preserved and faithfully transmitted through cell division. For genes that are silenced, heterochromatin domains enriched with specific histone post-translational modifications provide the structural and molecular basis for repression. H3K27 methylation marks facultative heterochromatin that is dynamically regulated across cell lineages and maintains the silencing of developmental genes^1,2^. As H3K27 methylation accumulates on chromatin, it forms Polycomb-silenced domains enriched for H3K27 di-methylation (H3K27me2) and tri-methylation (H3K27me3). This raises a fundamental question: how are the H3K27me2 and H3K27me3 marks propagated during DNA replication and chromosome duplication for heritable gene silencing^3,4^?

Polycomb Repressive Complex 2 (PRC2) in mammals consists of the core subunits EED, EZH2/1, RBAP46/48 and SUZ12, and catalyzes progressive methylation of the H3K27 residue. Catalysis by PRC2 proceeds rapidly from the unmethylated residue to H3K27me1 and then H3K27me2, however the conversion to H3K27me3 occurs with a 10-fold reduced catalytic rate^5,6^. This final methylation step is enhanced by allosteric activation of PRC2 when the EED subunit binds to a H3K27me3 mark on a neighboring nucleosome^5,7^. PRC2 allosteric activation catalyzes spreading of the H3K27me3 mark along chromatin fibers to form Polycomb domains^8,9^. However, the intermediate H3K27me2 modification is much more abundant, occupying 50-70% of histone H3 tails in chromatin^5,7,10^. The abundance of the H3K27me2 modification suggests that widespread PRC2 activity occurs without DNA sequence-targeting.

How are these methylation patterns in the genome propagated through chromatin duplication and cell division? In plants, the ATXR5/6 methyltransferases associate with the DNA replication fork and catalyze H3K27 methylation on newly deposited histones. This restores tri-methylation patterns by the end of S-phase^11^. In mammals, the EZH2 methyltransferase subunit of PRC2 co-precipitates with the replisome subunit PCNA and the chromatin assembly factor 1 (CAF1) that deposits new histones after the DNA replication fork^12^. Studies in mammals suggest that PRC2 catalysis at replication forks^9^ maintains the H3K27me3 mark by propagating it onto new, unmodified histones. However, this model has been questioned by the slow catalysis of the mark, which is not recovered during S-phase after DNA replication^13–18^.

In this study we use CUT&Tag chromatin profiling to track the dynamics of H3K27 methylation states during the cell cycle in human K562 cells. Comparison of H3K27 methylation dynamics to replication timing, the H3K27ac modification, and active RNA polymerase II (RNAPII) reveals that Polycomb-silenced domains are transiently marked by the H3K27me1 and H3K27me2 modifications as stepwise methylation restores H3K27me3 patterns after DNA replication. Outside of Polycomb domains, thousands of active genes gain the H3K27me1 mark after DNA replication, while genes with weak promoter activity proceed to H3K27me2. We find that sequential catalysis of H3K27 methylation is slow at genes that replicate in early S-phase, and fast later in S-phase. These replicative patterns of H3K27 mono– and di-methylation are catalyzed by PRC2, as they are lost upon either catalytic or allosteric inhibition of the complex. Upon inhibition, chromatin accumulates H3K27 acetylation. Finally, we show that the H3K27me2 mark does not repress transcription, which suggests a passive process at inactive genes. Together, our results explain how H3K27 modification patterns are faithfully duplicated in proliferating cells.

## Results

### The steady-state distributions of histone H3K27 modifications in K562 cells

To determine a baseline for analyzing the cell cycle dynamics of the three H3K27 methylation states, we characterized their steady-state distributions in the human erythroleukemic K562 cell line. We performed profiling of mono-, di-, and tri-methylation via CUT&Tag^19^ chromatin profiling using highly specific antibodies validated on synthetic nucleosomes^20^ (Supplementary Fig. 1). To identify active genes, we also profiled histone H3K27 acetylation (H3K27ac), and RNA Polymerase II (RNAPII) phosphorylated at serine 2 and/or 5 (active RNAPII). As expected, large >10-kilobase (kb) regions are enriched for the H3K27me3 modification, consistent with the annotation of Polycomb domains in K562 cells^21,22^ (Fig. 1A, Supplementary Fig. 2). For example, a genomic region on Chromosome *6* shows an H3K27me3-marked Polycomb domain encompassing four genes next to two active genes (Fig. 1A). In this region, the Polycomb domain lacks H3K27me1 and H3K27me2 modifications, while the two active genes are marked with the H3K27me1 modification and little H3K27me2 modification. In contrast, the intervening regions are enriched with the H3K27me2 mark. Thus, these three methylation states of the histone H3 K27 residue mark largely distinct regions.

**Figure 1.**
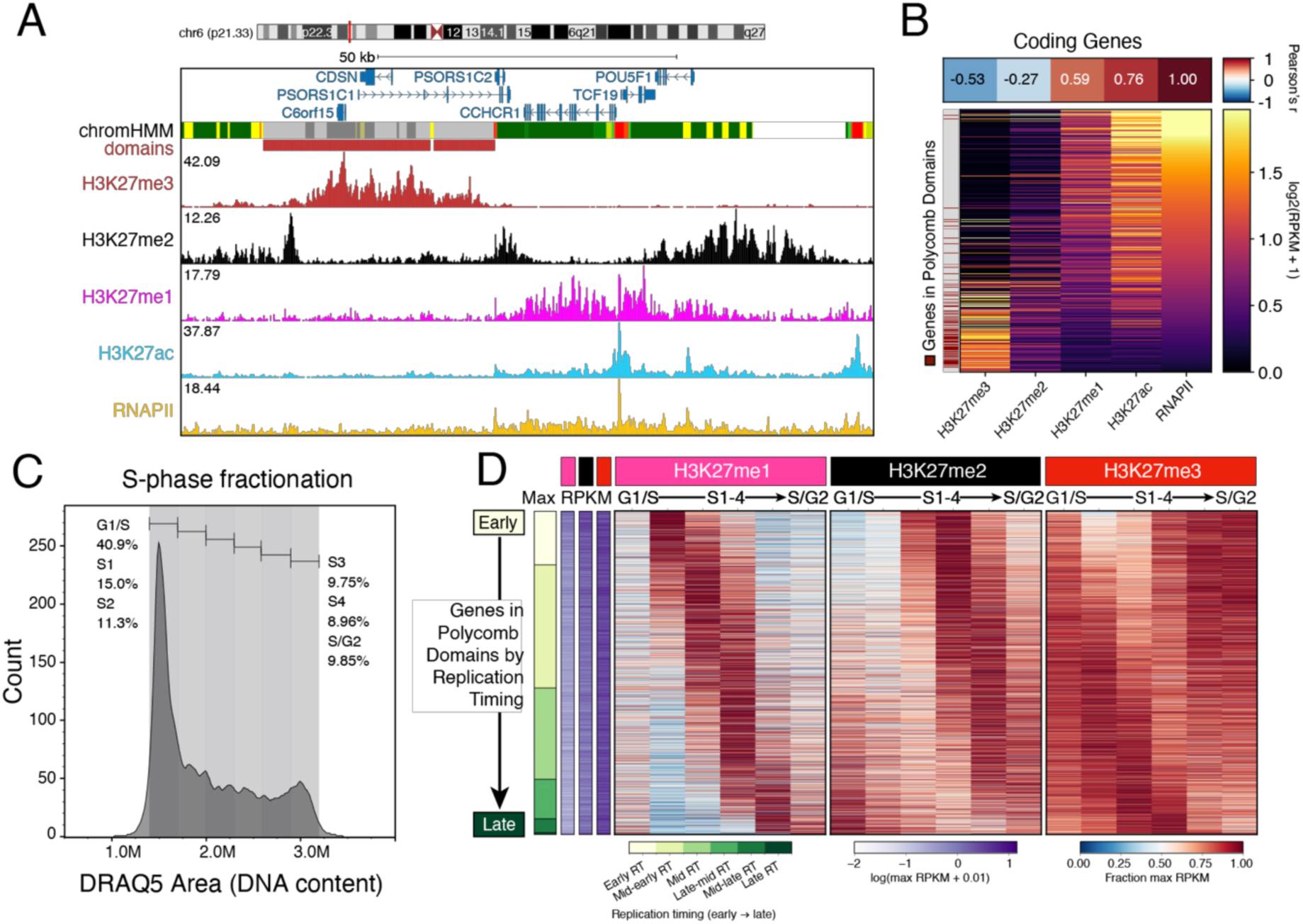
Stepwise methylation of histone H3K27 in Polycomb domains after DNA replication: (A) Browser snapshot of CUT&Tag profiling in K562 cells for the H3K27me3, H3K27me2, H3K27me1, and H3K27ac modifications, and RNAPIIS2/5p at the POU5F1 (OCT4) locus. A Polycomb domain^21^ overlaps the PSOR1S1C gene. (B) (top) Pearson correlation heatmap of active RNAPII versus H3K27 modifications at coding genes including a 1 kilobase promoter region upstream (n=19,946). (bottom) Signal heatmap of correlated gene values after log2 transformation. Genes in annotated Polycomb domains are highlighted in red and show high H3K27me3, low active RNAPII signal. (C) Gating schema based on DRAQ5 measurements of DNA-content used to isolate S-phase fractions via fluorescence-activated cell sorting (FACS). (D) CUT&Tag signal at genes in Polycomb domains (n=2,861) ranked by replication timing (y-axis) over the S-phase fractions (x-axis) from (C). Signal is shown as fraction of the maximum averaged across two replicates. Log10(max RPKM) per gene is shown in purple for the H3K27me1, H3K27me2, and H3K27me3 modifications to demonstrate high H3K27me3 signal and relatively less H3K27me2 and H3K27me1 signal at these genes.

To determine the relationship of each H3K27 modification with active RNAPII, we displayed their average signals at protein-coding genes (Fig. 1B). As expected, genes in Polycomb domains exhibited low active RNAPII. Similarly, genes enriched for the H3K27me2 mark are characterized by low RNAPII signal. In contrast, the H3K27me1 and H3K27ac modifications are enriched at active genes^23^. Our finding that di– and tri-methylated H3K27 vary inversely with active RNAPII at genes is consistent with previous observations in mouse embryonic stem cells^24,25^.

### Stepwise methylation of the histone H3K27 residue in Polycomb domains

For histone modifications to be stably maintained in the epigenome they must be duplicated every cell cycle after DNA replication. Chromatin duplication occurs via sequential events: (1) old, modified histones are distributively segregated to daughter strands, (2) new, unmodified histones are deposited on the strands to complete chromatin assembly, and (3) enzymatic catalysis modifies the new histones. Studies suggest that H3K27 tri-methylation occurs slowly after DNA replication^9,13–17^, but catalysis of the H3K27me1 and H3K27me2 marks during the cell cycle have not been examined. We therefore set out to resolve the cell cycle dynamics of all three methylation states in K562 cells.

We first asked how the steady-state levels of the H3K27 modifications correlate with transcription and timing in S-phase when a locus replicates. To compare replication timing with H3K27 modifications and active RNAPII, we visualized the density of 10-kb windows enriched for each mark along a continuous axis from early– to late-S-phase (Supplementary Fig. 3A-B). Windows enriched for H3K27me3 mostly replicated in mid-S-phase, although there are both earlier– and later-replicating windows. In contrast, H3K27me1-enriched windows predominantly replicated at the beginning of S-phase, as did windows enriched for H3K27ac and active RNAPII. Finally, H3K27me2-enriched windows replicated throughout S-phase. Thus, regions enriched for the H3K27me3, H3K27me2, and H3K27me1 marks exhibited distinct replication timing (Supplementary Fig. 3B). Our findings are consistent with the association of early replication timing and transcriptional activity^26^.

To separate cells during S-phase for CUT&Tag chromatin profiling of H3K27 methylation, we isolated 6 fractions of cells by fluorescence-activated cell sorting (FACS) on DNA content, where each fraction represents about 2.5-3 hours of the cell cycle (Fig. 1C). We then profiled each H3K27 residue methylation, the H3K27ac modification, and active RNAPII in each fraction. We first analyzed the 2,855 genes in Polycomb domains because new histones deposited there after DNA replication must become tri-methylated. We ranked genes based on their replication timing^27^, assigned them to six bins from early to late S-phase, and then displayed their chromatin profiling signals in FACS-isolated fractions (Fig.1D). We expected genes in Polycomb domains to undergo two-fold dilution of the H3K27me3 mark via distributive histone segregation, and then recover the mark as PRC2 methylates new histones. Indeed, early-replicating Polycomb domains have their lowest H3K27me3 signals in S1 and S2 fractions, while late-replicating Polycomb domains have their lowest signals in S4 and S/G2 fractions. This low methylation persists for 1-2 fractions after replication, indicating a delay of at least ∼5-6 hours before new histones are tri-methylated. This delay appeared consistent regardless of when genes in Polycomb domains replicate, with early-replicating Polycomb domain genes starting to recover tri-methylation by late S-phase and late-replicating domains beginning tri-methylation in the next cell cycle (Supplementary Fig. 4).

Polycomb domains have low mono– and di-methylation in steady-state populations (Fig. 1A). Strikingly, these marks transiently increase during S-phase in FACS-isolated fractions (Fig. 1D). Mono-methylation appears as a transient wave just as genes replicate and then rapidly diminishes (Fig. 1D, Supplementary Fig. 4). Di-methylation appears slightly later by 1-2 fractions after the mono-methylation wave and then diminishes as well. These patterns are simply explained by distributive catalysis by PRC2 in Polycomb domains, where: (1) mono-methylation occurs rapidly on new histones behind the replication fork, (2) mono-methylation is converted to di-methylation, and (3) eventually, di-methylation is converted to tri-methylation.

### Stepwise methylation at genes outside Polycomb domains

In steady-state populations, the H3K27me2 modification occurs outside Polycomb domains (Fig. 1A-B), so we next set out to analyze the H3K27 methylation dynamics in these regions. Genes outside Polycomb domains were categorized based on the most abundant H3K27 modification above a minimum signal threshold (Supplementary Table 1). This classification schema identified 2,268 H3K27me2-, and 3,872 H3K27me1-enriched genes (Supplementary Fig. 5A-B). These H3K27me2– and H3K27me1-enriched genes are not targets of PRC2 or PRC1 subunits SUZ12, CBX2 and CBX8 (Supplementary Fig. 5C), suggesting that they result from untargeted catalysis. The H3K27me1 genes were enriched for MYC, RNAPII, EP300, and other general transcription factors (Sup Fig. 4C). In contrast, H3K27me2-enriched genes were not significantly enriched for any of the 116 factors profiled in K562 cells by the ENCODE consortium, suggesting that they are inactive^21,28^.

We hypothesized that stepwise catalysis also occurs outside domains but does not achieve tri-methylation. To test this hypothesis, we displayed the signal of each mark at H3K27me2-enriched genes outside domains and grouped them by replication timing (Fig. 2A). The stepwise gain of mono– and di-methylation was recapitulated at genes enriched for H3K27me2 (Fig. 2A). In general, tri-methylation did not appear to follow from di-methylation at H3K27me2 genes as in Polycomb domains (Fig. 2A, Supplementary Fig. 6). The H3K27me1-enriched genes also gained mono-methylation, but it was not converted to di-methylation in the next fraction (Fig. 2B). Therefore, mono– and di-methylation of the H3K27 residue occurs after DNA replication at thousands of genes outside of Polycomb domains.

**Figure 2.**
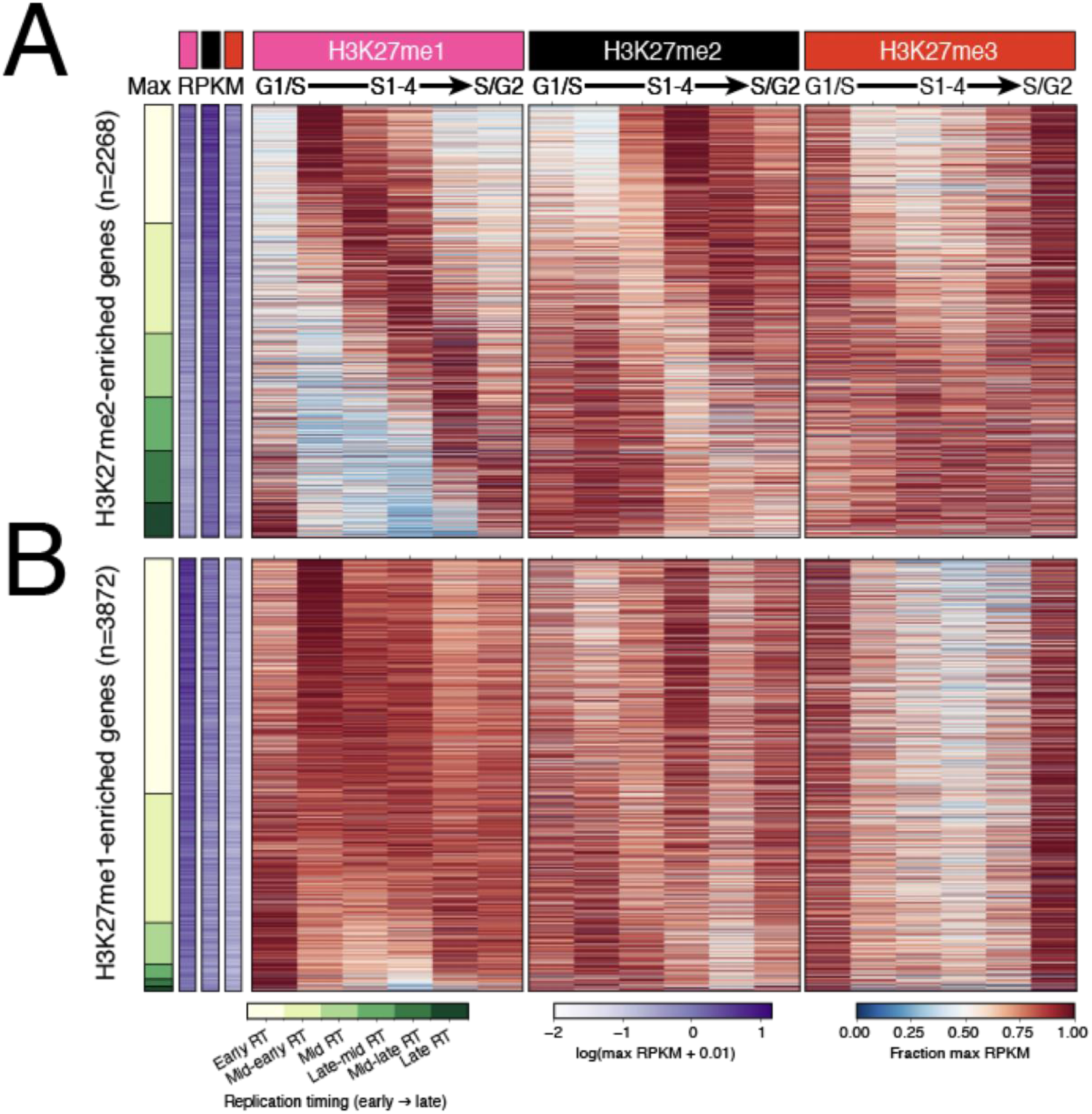
Stepwise methylation of the H3K27 residue outside of Polycomb domains: (A) CUT&Tag signal at genes outside Polycomb domains enriched for H3K27me2 (top, n=2,268) and (B) H3K27me1 (bottom, n=3,872) ranked by replication timing (y-axis) over the S-phase fractions (x-axis). Signal is shown as fraction of the maximum (dilution) averaged across two replicates.

### Increased catalysis of H3K27 methylation in late S-phase

To compare the dynamics of H3K27 methylation at early– versus late-replicating genes, we grouped Polycomb domain and H3K27me2-enriched genes by replication timing and visualized the complete cycle of H3K27 methylation as the magnitude of change (ΔRPKM) between S-phase fractions (Fig. 3). To resolve conversion of the H3K27 methylation states, we compared the change between marks (Fig. 3A). For example, a clockwise cycle indicates conversion of mark X (x-axis) to mark Y (y-axis). Compression of the x-axis only indicates fast conversion of mark X to mark Y. Compression of the y-axis indicates less catalysis of mark Y, and compression of both axes indicates less change in both marks overall. Lastly, a counterclockwise cycle indicates conversion of mark Y to mark X. We observed each of these by comparing the H3K27me2 mark to H3K27me3 and H3K27me1 (Fig. 3B-C, top and bottom row, respectively).

**Figure 3.**
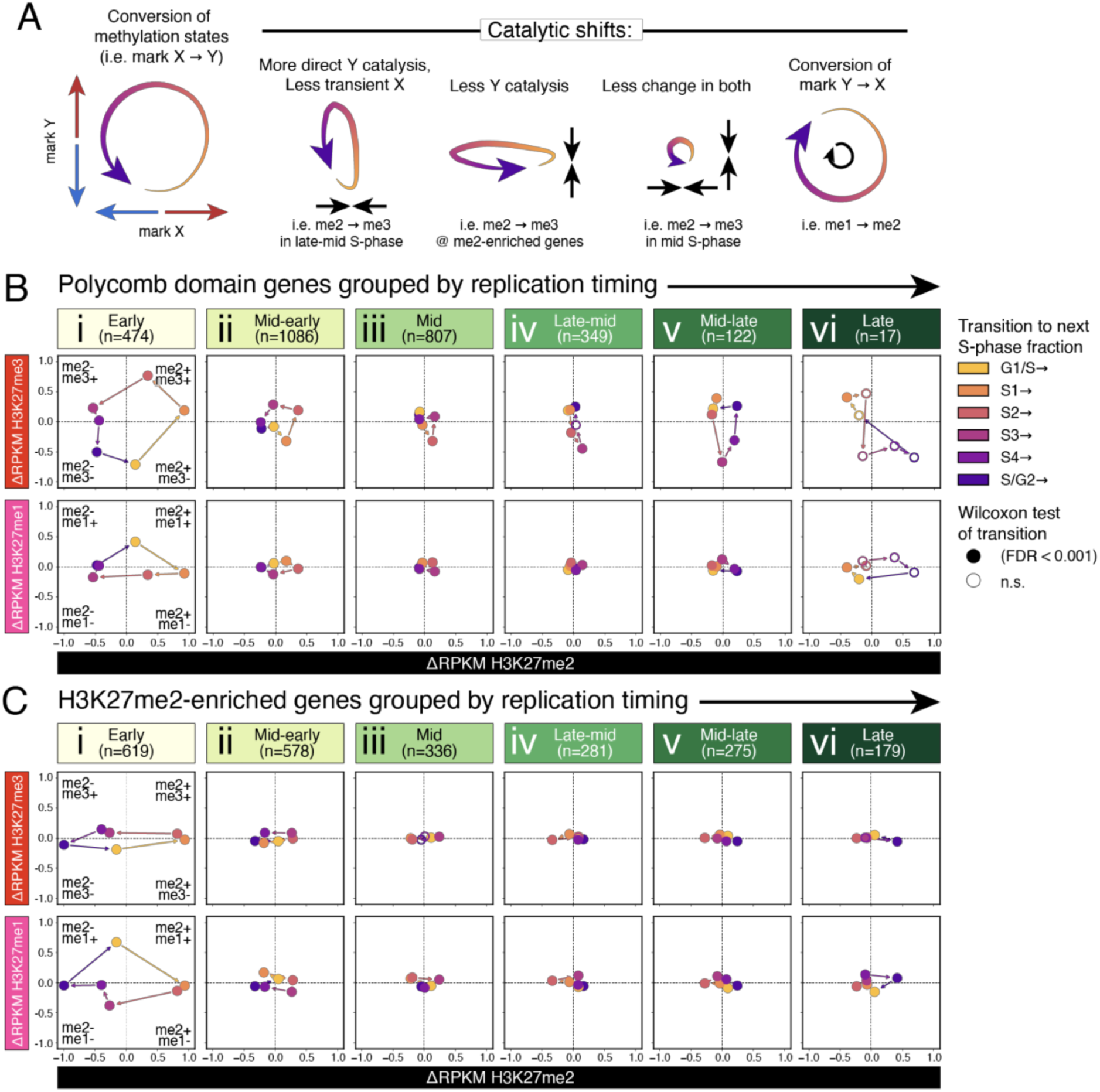
A cycle of H3K27 methylation at genes inside and outside of Polycomb domains: (A) diagram of possible catalytic patterns. (B) magnitude of change shown as DRPKM between S-phase fractions for H3K27me2 versus H3K27me3 (top) and H3K27me1 (bottom) for genes in Polycomb domains grouped by replication timing. Filled dots are medians with a significant change in either mark. Arrows show the direction of the cell cycle time course between S-phase fractions. (C) Same as (B) but for H3K27me2-enriched genes.

At Polycomb domain genes that replicate in early S-phase (Fig. 3B i), we observed well-defined changes in H3K27 methylation catalysis between the G1/S and the S1 fractions (Fig. 3, yellow). During this interval in aggregate, tri-methylation decreased by –0.7 ΔRPKM while mono– and di-methylation increased by +0.5 and +0.15 ΔRPKM, respectively. This was followed after the S1 fraction (Fig. 3, orange) by the loss of mono-methylation (–0.2 ΔRPKM) and gain of large amounts of di-methylation (+1 ΔRPKM) which began to convert to tri-methylation (+0.15 ΔRPKM). Tri-methylation then increased by +0.8 ΔRPKM while di-methylation dropped by 0.5 ΔRPKM from the S2 to the S3 fraction (Fig. 3, peach). Mono-methylation continued to decrease by –0.2 ΔRPKM from S3 to the S4 fraction (Fig. 3, magenta) while di-methylation also decreased (–0.5 ΔRPKM) and tri-methylation increased by +0.2 ΔRPKM. Catalysis of tri-methylation then ceased after the S4 fraction (Fig. 3, light purple). Both di– and tri-methylation rapidly decreased by –0.5 ΔRPKM from the S/G2 to the G1/S fraction (Fig. 3, dark purple), thus forming a complete cycle, consistent with stepwise catalysis.

Notably, early replicating genes in Polycomb domains (Fig. 3B i) exhibited significantly greater loss and gain in H3K27 tri– and in di-methylation than genes that replicated later in S-phase (Supplementary Fig. 7a i-ii). At late-mid replicating genes (Fig. 3B iv) H3K27me1 dynamics were not apparent. These changes suggest that the catalysis of H3K27me3 is slow in early S-phase but rapid in late S-phase. In general, this shift toward H3K27me3 catalysis coincided with S-phase progression as the change in all marks diminished (Fig. 3B, Supplementary Fig. 7A). Our analysis was inconclusive for late-replicating genes (Fig. 3B vi) due to low sample size.

At the H3K27me2-enriched genes outside Polycomb domains, we were able to distinguish the conversion of H3K27 mono– to di-methylation (Fig. 3C). These two marks formed a dynamic cycle as in Polycomb domains but conversion of H3K27 di– to tri-methylation was limited. As in Polycomb domains, we found a drop in the overall change of H3K27me2 and other marks after early S-phase (Supplementary Fig. 7B). We conclude that overall H3K27 methylation proceeds slowly in early S-phase and speeds up in late S-phase.

### PRC2 inhibition slows H3K27 methylation in the cell cycle

If stepwise catalysis is responsible for the transient appearance of H3K27me1 and H3K27me2 marks in Polycomb domains, then chemical inhibition of the EZH2/1 methyltransferase subunit of PRC2 should ablate these marks during the cell cycle. To test this, we treated cells for 8 hours with small molecule inhibitors of PRC2 followed by S-phase fractionation and CUT&Tag profiling (Fig. 4A). Since the cell cycle of K562 cells is ∼20 hours^29^ and fractions are isolated from the same parent population, we can detect temporal drug effects by comparing count-normalized changes between early– and late-replicating genes in fractions. For example, in the S1 fraction only early-replicating genes acquire new histones in the presence of the drug. Only these genes should be affected if a mark is catalyzed in step with DNA replication and those genes should be unaffected in later fractions.

**Figure 4.**
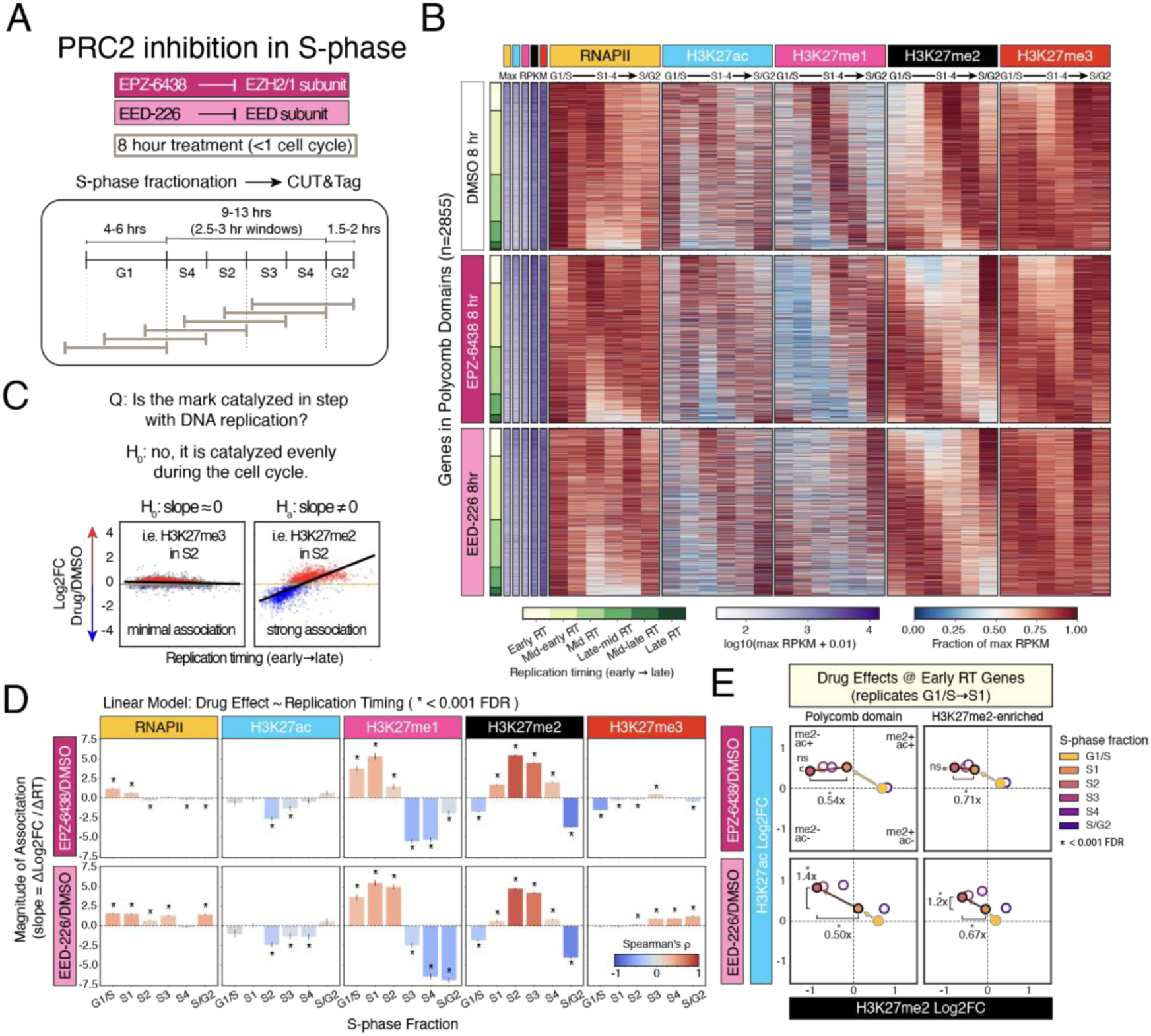
PRC2 inhibition delays H3K27me1 and H3K27me2 modifications in Polycomb domains after DNA replication: (A) Experimental design for 8-hour treatment of cycling K562 cells with small-molecule inhibitors of PRC2. (B) CUT&Tag signal at genes in Polycomb domains (n=2,861) ranked by replication timing (y-axis) over the S-phase fractions (x-axis). Signal is shown as fraction of the maximum averaged across two replicates. Max RPKM is computed per treatment across the S-phase time course. Log10(max RPKM) per gene is show in purple for active RNAPII, H3K27ac, H3K27me1, H3K27me2, and H3K27me3 (left to right). The heatmaps show signal after treatment for 8 hours with DMSO (vehicle control, top), the EZH2 inhibitor EPZ-6438 (middle), and the EED inhibitor EED-226 (bottom). (C) Statistical paradigm for testing the relationship between drug effects versus replication timing at genes in Polycomb domains. Scatter plots are from Supplementary Figure 8. Genes above an absolute of log2 fold-change of 0.2 are colored red for up in treatment or blue for down in treatment, or grey for Log2 fold-change < 0.2. (D) Linear regression analysis (y-axis) of response to EPZ-6438 (top), and EED-226 (bottom) within S-phase fractions (x-axis) versus replication timing at genes in Polycomb domains. Stars indicate a significance < 0.001 after FDR correction. Bar color is Spearman’s rho (E) PRC2 inhibition at early replicating genes in and outside Polycomb domains. Median log2 fold-change for EPZ-6438 (top) and EED-226 (bottom) versus DMSO is shown for H3K27me2 (y-axis) and H3K27ac (x-axis) in the S2 fraction. The fold difference between Log2 fold-change in the S1 versus S2 fraction is shown for differences greater than ±0.2. Stars indicate a significant difference in Log2 fold-change values by the Wilcoxon test with a p-value < 0.001 after Bonferroni correction.

We treated cells with either EPZ-6438, a catalytic inhibitor of the EZH2/1 methyltransferase subunit^30^, or with EED-226, which binds the EED subunit responsible for allosteric activation of the complex^31^. We observed a striking delay of ∼4 fractions for the H3K27me1 and H3K27me2 marks in Polycomb domains in both treatment conditions compared to the stepwise catalysis in the DMSO control (Fig. 4B). In contrast, H3K27me3 catalysis appeared far less affected as measured by the maximum RPKM per gene and signal trend across the time course. Due to count normalization, this could result from inhibited global catalysis irrespective of when a locus replicates.

We analyzed genes in Polycomb domains and compared the effects of the two PRC2 inhibitors normalized to the DMSO control versus replication timing (Fig. 4C). The change in each S-phase fraction reflects the acute effects of inhibitors on H3K27 methylation for 8 hours preceding sample collection. We therefore hypothesized that H3K27 methylation changes in S-phase fractions will linearly relate to replicating timing if methylation follows in step with DNA replication (Fig. 4C). The magnitude of association would be revealed in the slope of a simple linear model, where a value of zero indicates no association and a positive slope indicates early-replicating genes are relatively depleted for the mark, while a negative slope indicates late-replicating genes are depleted.

The effects of the two inhibitors were generally consistent across the H3K27 mono– and di-methyl marks, and genes were depleted of the marks during replication in the presence of the drugs (Fig. 4D, Supplementary Fig. 8, Supplementary Table 2). In contrast, the inhibitors had little effect on the tri-methyl mark (Fig. 4B,D). We therefore conclude that the transient H3K27 mono– and di-methylation following replication are acutely affected by both catalytic and allosteric inhibition of PRC2. We attribute the lack of effect on H3K27 tri-methylation to the slow catalysis of this mark, such that it is not achieved in the untreated cells in 8 hours.

We also found consistent effects of both drugs on active RNAPII and the H3K27ac mark at genes in Polycomb domains (Fig. 4B). Our analysis allowed us to temporally link changes in H3K27 methylation with shifts in H3K27ac and active RNAPII signals. The linear relationship between replication timing and the inhibitors’ effects on active RNAPII were close-to-zero; these changes appear unrelated to replication timing (Fig. 4D). In contrast, the relationship with the H3K27ac mark was negative for both inhibitors, but only in the S2 and S3 fractions (Fig. 4C, Supplementary Fig. 8). At these time points, many genes will have replicated in early S-phase in the presence of the PRC2 inhibitors

To resolve exactly when the changes in the H3K27ac and H3K27me2 marks occurred at early replicating genes in Polycomb domains, we plotted the change in one mark versus the other in each fraction (Fig. 4E, left). The time when the H3K27ac mark increased was not consistent across the two inhibitor treatments. Under EPZ-6348 treatment, no significant gain of the H3K27ac mark occurred during the interval from the S1 to the S2 fraction (Fig. 4E, upper left, Supplementary Fig. 9A i). Rather, the gain in acetylation occurred one fraction earlier during the interval from the G1/S to the S1 fraction before the H3K27me2 mark was lost. Under EED-226 treatment, an increase of H3K27ac by 1.4x occurred in step with the loss of the H3K27me2 mark (Fig. 4E, bottom left). To control for any loss of the H3K27me3 mark in Polycomb domains, we analyzed H3K27me2-enriched genes outside Polycomb domains and observed similar timing (Fig. 4E, right column). These patterns were clear at early replicating genes, but not at later replicating genes (Supplementary Table 3). We therefore conclude that catalytic inhibition of PRC2 indirectly allows the acetylation of the H3K27 residue at early replicating genes, and allosteric inhibition does so more slowly.

### H3K27me2-marked regions are inactive

We next sought to define the functional relationship between the H3K27me2 mark and gene transcription. At steady-state, this methylation mark is anticorrelated with active RNAPII in genes (Fig. 1B), so we asked whether the mark is repressive *per se* by querying promoter activity in three contexts in K562 cells (Fig. 5A). First, as a measure of promoter activity in the native chromatin context, we used data obtained with the GRO-cap (global run-on sequencing with 5′ cap selection) method^32^ (Fig. 5B). Second, to measure autonomous promoter activity when the DNA sequence is moved from native context to a plasmid reporter, we used data obtained with the SuRE (Survey of Regulatory Elements) method^33^ (Fig. 5C). Lastly, to determine whether the H3K27me2 mark represses transcription, we asked whether promoter sequences are repressed when integrated into regions marked by H3K27me2 using the TRIP (Thousands of Reporters Integrated in Parallel) method^34,35^ (Fig. 5D). In the native context, promoters of H3K27me2-marked genes had weak median activity that was 9-fold higher than those of Polycomb domain genes (Fig. 5B). In contrast, promoters of H3K27me1– and H3K27ac-marked genes had 699-fold and 1437-fold higher median native activity than those of Polycomb domain genes, respectively. The difference between promoters of Polycomb domain genes and H3K27me2-marked genes was erased when these promoters were moved to a plasmid. In contrast, the promoters of H3K27me1– and H3K27ac-marked genes had 16-fold and 21-fold higher median autonomous activity than those of Polycomb domain genes (Fig. 5C). In summary, we found that promoters of genes marked by H3K27me2 are relatively inactive both in their native context and when moved to a plasmid.

**Figure 5.**
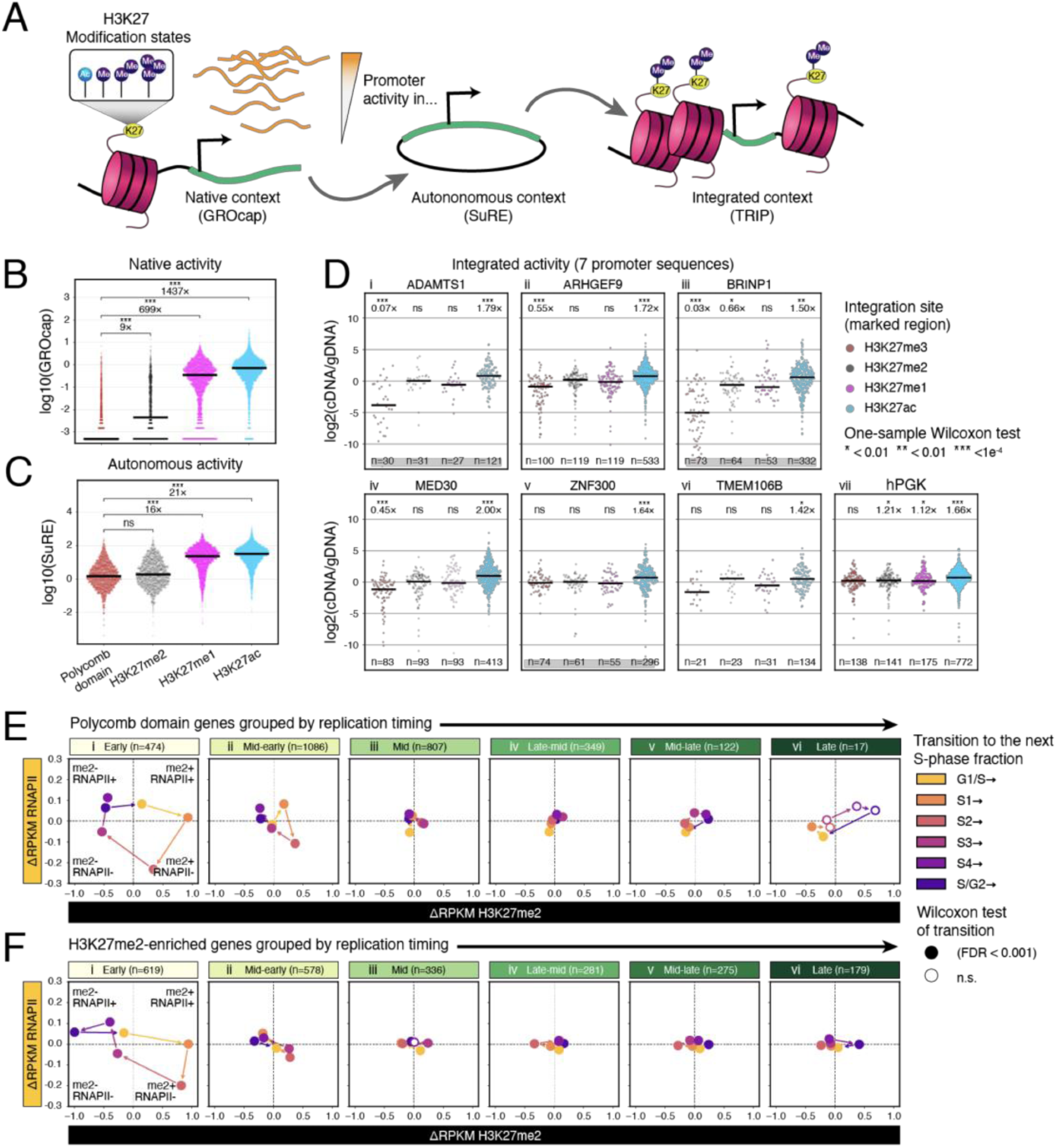
Histone H3K27me2 modification marks inactive chromatin: (A) Schematic of experimental design to measure promoter activity in three different contexts. (B) Native promoter activity measured by GROcap^32^ signal at gene promoters grouped by most abundant H3K27 modification (defined in Supplementary Fig. 5A). (C) Same as (B) but for SuRE^33^ signal. (D) Expression of promoters integrated into H3K27-modified chromatin. TRIP data are from Ref.^34^. Dot plots are expression values per integration normalized to genomic DNA with median per group. H3K27 classifications are the top 5% of regions defined in Supplementary Figure 9E. FDR-corrected one-sample Wilcoxon test (null: log2(cDNA/gDNA = 0) and fold change is computed against that expectation (E) magnitude of change shown as delta-RPKM between S-phase fractions for active RNAPII versus H3K27me2 at genes in Polycomb domains grouped by replication timing. Filled dots are medians with a significant change in either mark. Insignificant change versus the null of no change is denoted by no fill. Arrows show the direction of the cell cycle time course between S-phase fractions. (F) Same as (E) but for H3K27me2-enriched genes.

We next asked whether seven promoter sequences^34^ are repressed when integrated into the regions uniquely marked by H3K27me2 (Fig. 5D, Supplementary Fig. 10). Three of these promoters are repressed in their native chromatin context (*ADAMTS1, ARHGEF9*, and *BRINP1)* three are weakly expressed (*MED30, ZNF300*, and *TMEM106B)*, and one is constitutively expressed (*hPGK*). When integrated into the regions marked only by the H3K27me2 modification, one of these promoters (Fig. 5D iii) was significantly repressed, one was not repressed (Fig. 5D vi), and five were neutrally affected. H3K27me1-marked regions were also found to have a neutral effect on six out of seven promoters, and H3K27me3-marked regions significantly repressed four of the seven promoters. We therefore conclude that the H3K27me2 mark does not repress most gene promoters.

If the H3K27me2 mark is not repressive, then the slow dynamics of PRC2 might create a window for gene activation after DNA replication during the 1-2 fractions before tri-methylation is recovered. To compare cell cycle changes of the H3K27me2 mark versus active RNAPII at Polycomb domain genes, we visualized the magnitude of change (ΔRPKM) between S-phase fractions (Fig. 5E). At early replicating genes, which include many rapid stimulus response genes like *IL1B*, *IL17C*, and *IL10* that regulate inflammatory response (Supplementary Figure 11, Supplementary Table 1), the dynamics of these two marks formed a counter-clockwise cycle where modest gains in active RNAPII (<+0.1RPKM per fraction) preceded catalysis of di-methylation. The increase of active RNAPII initiated after S3 through the S4 fraction. This continued through DNA replication after G1/S, into S1 and until di-methylation increased in the S2 fraction (Fig. 5E i). The cycle compressed along both axes at mid-early replicating genes and was not discernable at genes replicating later in S-phase (Fig. 5E ii-vi). These include the homeotic patterning genes at the *HOXA* and *HOXD* loci (Supplementary Table 1) that are stably repressed in K562 cells. We thus found that early replicating genes in Polycomb domains transiently gain active RNAPII, before di-methylation is catalyzed, and significantly more so than later replicating genes (Supplementary Fig. 7A-B).

At the H3K27me2-enriched genes outside Polycomb domains, transient gains of active RNAPII were also greatest at early replicating genes (Fig. 5F i). These include other stimulus response genes like *IL11, TNF,* and *CCL24* (Supplementary Figure 12, Supplementary Table 1). Active RNAPII at these sites increased only when H3K27me2 decreased or did not change, such that changes in the two marks were inversely related, both at early and mid-early replicating genes (Fig. 5F i-ii). This reveals that loading of RNAPII and catalysis to H3K27 di-methylation are incompatible.

## Discussion

Previous studies dissecting the dynamics of chromatin duplication have shown that the H3K27me3 modification, which is essential for developmental gene silencing, can take as much as 24 hours to regain pre-replication levels^14,15^. The slow recovery kinetics of the H3K27me3 mark challenge the model where it is propagated onto new, unmodified histones by PRC2 allosteric activation at replication forks^9^. Here, we resolve the catalysis of H3K27 methylation after DNA replication in cycling K562 cells and find that tri-methylation at genes in Polycomb domains proceeds via sequential mono– and di-methyl intermediates in the timespan of hours per step. This sequential catalysis in Polycomb domains proceeds to tri-methylation via allosteric activation of PRC2^5,7,8^. In active genes histone methylation stops at H3K27me1. This may be due to transcription of DNA that disengages the PRC2 complex, turns over nucleosomes^36,37^, or recruits the UTX H3K27 demethylase^38,39^. In contrast, at inactive genes and other sites of low transcription, sequential catalysis yields the H3K27me2 mark. The H3K27me2 modification is one of the most abundant histone modifications across metazoan genomes, and has been proposed to limit pervasive transcription^40^. However, we find no evidence that the H3K27me2 mark is associated with transcriptional silencing in K562 cells. We conclude that the H3K27me2 modification is not repressive, but rather simply accumulates on inactive chromatin.

PRC2 enzymatic activity is understood based on the biochemical kinetics of the complex on *in vitro* assembled nucleosome substrates^5,41^. *In vitro*, the presence of tri-methylated H3K27 peptides greatly enhances efficient catalysis of H3K27me3 on an unmodified nucleosome substrate via allosteric activation of PRC2 at the EED subunit^5^. The subsequent observation of PRC2 at replication forks led to the model where PRC2 allosteric activation there reads H3K27me3 on old histones and writes H3K27me3 on new ones^9^. Structural studies have also documented PRC2 complexes with other allosteric activators that are suggested to promote processive methylation of the unmodified H3K27 residue to H3K27me3^7,42^. We find using chromatin profiling in S-phase fractions that H3K27 methylation occurs in sequential steps via mono-, di-, and tri-methylation of the residue *in vivo*. These events can occur over the course of hours, which supports largely distributive methylation by PRC2 where each binding event adds one methyl group.

Chromatin profiling of S-phase fractions provides a rough timing of chromatin dynamics over the course of hours. In K562 cells the doubling time is ∼20 hours: G1-phase is 4-6 hours, S-phase is 9-13 hours, and G2-phase is 1.5-2 hours^29^. We find that H3K27me3 is fully regained over the course of 2-4 fractions. Changes in the H3K27 methyl marks are greatest at early replicating genes in and outside Polycomb domains. After DNA replication later in S-phase, stepwise H3K27 methylation is enhanced to promote more efficient catalysis of H3K27me3. That early-versus later-replicating genes experience different rates of PRC2 catalysis may result from increased allosteric activation of the complex later in S-phase. This could result from the limited decompaction of replicating Polycomb domains^43^ which maintains a high density of H3K27me3-marked nucleosomes that thereby enhance PRC2 allosteric activation. Many cellular factors change during S-phase progression, and our data suggests that proximity to the onset of DNA replication^44,45^ after G1-phase shapes PRC2 kinetics during S-phase.

### The H3K27me2 modification accumulates after DNA replication at inactive chromatin

In embryonic development H3K27 methylation is essential for gastrulation^2^ and lineage acquisition^46,47^. PRC2 first catalyzes the H3K27me2 mark at sites with histone H2A ubiquitination after zygotic genome activation in flies, zebrafish, and mice^48–50^. These sites later form Polycomb domains marked by H3K27me3. During embryogenesis nearly all H3K27me2 matures into H3K27me3, but pre-epiblast mouse embryonic stem cells (mESCs) have H3K27me1, H3K27me2, and H3K27me3 that is mutually distinct: H3K27me1 covers active gene bodies, H3K27me2 spans intergenic regions and inactive genes, and H3K27me3 is focused around CpG-rich sequences which occur at ∼70% of mammalian promoters^24,25^. Our data find that H3K27me2 results from widespread catalysis in S-phase which may not depend on pre-existing histone H2A ubiquitination.

During H3K27 methylation by PRC2, H3K27me2-modified nucleosomes are the substrate for the H3K27me3 mark. This might suggest that inactive sites marked by H3K27me2 are more likely to gain the H3K27me3 mark. The maintenance of H3K27me2-marked regions devoid of the H3K27me3 mark across cell cycles means this conversion is restricted by the slow kinetics of PRC2 and low local allosteric activation of the complex. This implies that cells with different cell cycle lengths may achieve very different global distributions of the H3K27me3 mark^16^.

### PRC2 opposes acetyltransferase activity

Whereas H3K27me3 marks Polycomb domains, H3K27 acetylation marks active regulatory elements. Hindrance of H3K27 acetylation is achieved by H3K27me3 and H3K27me2, but not by H3K27me1^51^. Previously di-methylated regulatory elements gain H3K27ac upon PRC2 knockout and are activated in the early mouse embryo, mESCs and *Drosophila* cells, consistent with the idea that di-methylation blocks acetylation^25,40,48^. Our data corroborates that H3K27 acetylation is gained upon PRC2 inhibition and resolves the timing of H3K27 acetylation with respect to changes in H3K27 di-methylation in each cell cycle.

We found that PRC2 catalytic and allosteric inhibitors affect H3K27 methylation in S-phase and allow the gain of H3K27ac. After DNA replication in early S-phase, both drug types result in H3K27ac gains, but there is a slightly faster effect of EPZ-6438, suggesting that catalytic inhibition somehow allows more acetylation. This informs development of PRC2 inhibitors and their application toward treating multiple cancers^30,31,52^, because it suggests targeting different parts of PRC2 can shape cancer response^53^. Catalytic PRC2 inhibitors, including EPZ-6438, mimic S-adenosyl methionine and occupy the catalytic site of EZH2/1 methyltransferase subunit to form a catalytic block. This likely prevents PRC2 from binding the H3K27 residue. In contrast, allosteric PRC2 inhibitors, including EED-226, bind the aromatic cage of the EED subunit and allosterically modulate the EZH2/1 catalytic site. If PRC2 in this context can still bind the H3K27 residue after DNA replication, then PRC2 itself could shield the residue from acetylation.

PRC2 catalytic and allosteric inhibitors are suggested to have distinct effects on catalysis of H3K27me3^17^. In K562 cells, both catalytic and allosteric inhibition slow catalysis of H3K27me1 and H3K27me2 after DNA replication. In this way, PRC2 allosteric activation modulates not only catalysis of H3K27me3, but catalysis of all H3K27 methyl marks.

## Methods

### Cell culture

Human K562 cells (ATCC, Manassas, VA, Catalog #CCL-243) were cultured following the supplier’s protocol. For inhibitor treatments, stock solutions in DMSO were diluted in cell culture medium and added directly to the cultures at the specified final concentrations. The PRC2 inhibitors used were EPZ-6438 at 10 μM (Selleckchem, Catalog # S7128), EED-226 at 10 μM (Selleckchem, Catalog # S8496). DMSO at 1:1,000 v/v was used as control. Cells were harvested after 8 hours of treatment and processed for CUT&Tag profiling.

Drosophila S2 cells (RRID: CVCL_Z232) were grown to log phase in HYQ-SFX Insect medium (Invitrogen) supplemented with 18 mM L-glutamine prior to harvest. Cell counts and cell sizes were measured using the Vi-CELL XR Cell Viability Analyzer (www.beckman.com).

### Antibodies

We used the following antibodies: Guinea Pig anti-Rabbit IgG (Heavy & Light Chain) (Antibodies Online, Cat# ABIN101961), H3K27me3 (Rabbit monoclonal anti-H3K27me3, Cell Signaling Technology, Cat# 53207), H3K27me2 (Rabbit polyclonal anti-H3K27me2, Abcam, Cat# ab24684), H3K27me1 (H3K27me1 Recombinant Polyclonal Antibody, Thermo Fisher Scientific, Cat# 712817), H3K27ac (Rabbit monoclonal anti-H3K27me3, Cell Signaling Technology, Cat# 53207), RNAPIIS2/5P (Rabbit monoclonal Phospho-Rpb1 CTD (Ser2/Ser5), Cell Signaling Technology, Cat# 13546), H3K4me3 (Rabbit monoclonal anti-H3K4me3, EpiCypher, Cat# 13-0060), H3K4me2 (Rabbit monoclonal anti-H3K4me2, EpiCypher, Cat# 13-0027), and H3K4me1 (H3K4me1 Recombinant superclonal Antibody, Thermo Fisher Scientific, Cat# 710785). The final concentrations of the antibodies used in CUT&Tag are specified in the following sections. Antibodies were validated using the SNAP-CUTANA™ K-MetStat Panel according manufacturers guidelines (EpiCypher, Cat# 19-1002).

### Fluorescence-Activated Cell Sorting

To purify S-phase fractions of K562 cells, 10 million cells were resuspended in ice-cold harvest buffer (1x PBS with 2 mM EDTA, 2% FBS, 1 mM Spermidine, 0.05% Triton-X-100) and briefly fixed with 0.1% PFA for 2 minutes. Fixation was quenched with 2.5 M glycine for 5 minutes, then cells were spun down and resuspended in harvest buffer with 2.5 μM DRAQ5 (Thermo Scientific, 62251) to stain DNA. After at least 10 minutes of incubation, 6 cell cycle fractions were isolated on a BD Discover S8 imaging flow cytometer based on DRAQ5 intensity.

### CUT&Tag-direct for whole cells

CUT&Tag reactions were performed according to the CUT&Tag-direct protocol^54^ with minor modifications. 25,000 cells (no treatment) or 8,000 cells (DMSO, EPZ-6438, and EED22-6 treatment) were aliquoted in harvest buffer. *Drosophila* S2 cells were added to each sample at a ratio of 1.6:1. 2 μl of Bio-Mag Plus Concanavalin A (ConA)-coated magnetic beads (Bangs Laboratories Catalog #BP531) per sample were activated and added to the cells, then incubated for 10 min at room temperature. ConA-bound cells were suspended in wash buffer (20 mM HEPES pH 7.5, 150 mM NaCl, 0.5 mM spermidine, 0.05% Triton-X100, Roche Complete Protease Inhibitor EDTA-Free tablet, 2 mM EDTA) and split into individual 0.5 ml tubes for overnight incubation with the primary antibody at a 1:50 dilution at 4°C. After removing unbound primary antibody by washing, the samples were resuspended in wash buffer containing the secondary antibody (guinea pig anti-rabbit IgG) at a 1:100 dilution and incubated at room temperature for 1 h. Following another wash, the samples were resuspended in 300-wash buffer (wash buffer with an additional 150 mM NaCl) containing Protein AG-Tn5 (pAG-Tn5 at a 1:20 dilution, EpiCypher, Catalog #15-1117) and incubated at room temperature for 1 h. The samples were then washed in 300-wash buffer and resuspended in tagmentation buffer (300-wash buffer with 10 mM MgCl2), followed by incubation at 37°C for 1 h to complete the Tn5 tagmentation reaction. The samples were washed with TAPS wash buffer (10 mM TAPS with 0.2 mM EDTA) and resuspended in 5 μl of release solution (10 mM TAPS, 1% SDS, and 1:10 Thermolabile Proteinase K (New England Biolabs, Catalog #P8111S)). They were then incubated in a thermocycler with a heated lid at 37°C for 1 h and 58°C for 1 h to release fragments. To quench SDS, 15 μl of 6% Triton-X100 was added. PCR was performed by adding 2 μl each of barcoded 10 mM i5 and i7 primer solutions and 25 μl of premixed KAPA PCR Master Mix (10 μl of HiFi buffer, 1.5 μl of 10 mM dNTPs, 1 μl of KAPA HiFi polymerase, and 29.5 μl of H2O) (Roche, Catalog #07958846001). The following cycling conditions were used: Cycle 1: 58 °C for 5 min; Cycle 2: 72 °C for 5 min; Cycle 3: 98 °C for 30 s; Cycle 4: 98 °C for 10 s; Cycle 5: 60 °C for 10 s; Repeat Cycles 4-5 11 times; 72 °C for 1 min; Hold at 12 °C. CUT&Tag libraries were cleaned with HighPrep Paramagnetic bead-based post PCR clean-up reagent (MagBio, Catalog # AC-60500) at a 1.3:1 (vol/vol) ratio of beads to sample, quantified on the Agilent 4200 D1000 TapeStation (Agilent, Catalog #5067-5584) and pooled for sequencing.

### DNA sequencing data processing

The size distributions and molar concentration of libraries were determined using an Agilent 4200 TapeStation and libraries were mixed to achieve equal representation as desired aiming for a final concentration as recommended by the manufacturer. Paired-end 50×50 bp sequencing on the Illumina NextSeq 2000 platform was performed by the Fred Hutchinson Cancer Center Genomics Shared Resources and data were analyzed as described (https://www.protocols.io/view/cut-amp-tag-data-processing-and-analysis-tutorial-e6nvw93x7gmk/v1).

For processing sequencing data, we used cutadapt 4.4^55^ to trim adapters from 50 bp paired-end reads with parameters “-j 8 –-nextseq-trim 20 –m 20 –a

AGATCGGAAGAGCACACGTCTGAACTCCAGTCA –A

AGATCGGAAGAGCGTCGTGTAGGGAAAGAGTGT –Z”

We used Bowtie2 2.5.1^56^ to map the paired-end 50 bp reads to the hg38 human genome reference sequence from UCSC with parameters “--very-sensitive-local –-soft-clipped-unmapped-tlen –-dovetail –-no-mixed –-no-discordant –q –-phred33 –I 10 –X 1000”.

To remove spike-in reads, hg38 positions mapped to by reads from S2 profiles were blacklisted and reads that completely overlapped with of these intervals were removed using bedtools intersects. The spike-in reads were not used for data analysis.

### Data analysis and visualization

We used bedtools^57^ genomecov to generate bedGraph files, which were then converted to bigWig format using bedGraphToBigWig. Coverage-normalized bigWig files represent the fraction of counts at each base pair, scaled by the size of the reference hg38 genome (3,298,912,062), ensuring that if the counts were uniformly distributed, each position would have a value of 1. BigWigs were visualized in the UCSC genome browsers.

Reads overlapping the longest transcript of coding genes defined by gencode v48 plus one kilobase upstream were quantified using the Featurecounts function of the subread package. RPKM values per gene were calculated by dividing by total counts in millions and interval length in kilobases and then averaged across replicates.

### Replicating timing assignment

Processed replication timing data in K562 cells^27^ was obtained from the UCSC table browser for hg19 and lifted over to hg38 using CrossMap. Replication timing per gene was calculated as the average wavelet-transformed signal and inverted such that 0 is early and 1 is late. Six replication timing bins were fit to the distribution at even intervals between the 0.2 and 0.98 percentiles.

### Promoter activity quantification

Gene promoter regions were defined in^34^ and processed promoter values for GroCap and SuRE were obtained from the supplement. Processed TRIP data^34^ was also obtained from the supplement.

### H3K27 gene category assignment

Polycomb domains for 127 roadmap epigenomes^21^ were obtained from the WUSTL portal and defined as continuous ChromHMM intervals labelled “Repressed Polycomb”, “Weak Repressed PolyComb”, “Bivalent Enhancer”, “Flanking Bivalent TSS/Enh”, or “Bivalent/Poised TSS” greater than 1 kilobase. Genes were defined to be in Polycomb domains if they overlapped by at least 50%. The remaining genes were assigned to a H3K27 modification state based on the most abundant mark in S-phase fractions above a minimum threshold of 0.5 RPKM.

### Quantification and statistical analysis

For each CUT&Tag experiment, at least two biological replicates were included. Statistical comparisons between groups were performed using Pearson correlation and linear regression, as well as the Kolmogorov-Smirnov, Mann-Whitney, and Wilcoxon tests implemented in SciPy 1.17.0^58^. Bonferroni correction was applied to adjust for multiple comparisons and only adjusted p-values below the corrected threshold of 0.001 were considered statistically significant unless otherwise noted.

### Data and code availability

Raw data is available at GEO under accession XXX. Processed data is available at Zenodo under https://doi.org/10.5281/zenodo.19489947. Data processing and plotting scripts are publicly available on GitHub @ https://github.com/jacob-greene/H3K27me2. An interactive UCSC browser session with the manuscript data is available @ https://genome.ucsc.edu/s/jegreene/H3K27me2_manuscript.

## Acknowledgments

We thank Srinivas Ramachandran and Dominik Otto for valuable suggestions on data analysis and our Fred Hutchinson Cancer Center colleagues, including Jorja Henikoff for help with processing of sequencing data, Christine Codomo for sequencing library pooling, as well as Terri Bryson and Doris Xu for help with cell culture. This research was supported by the Howard Hughes Medical Institute (S.H.) and NIH grant no (Fred Hutchinson Cancer Center Shared Resources).

## Author contributions

J.E.G. performed the experiments and the analyses. K.A. and S.H. supervised the work. J.E.G. wrote the original draft. J.E.G., K.A., and S.H. reviewed and edited the draft.

## Supplemental Information

Classifications of protein-coding genes from Gencode v49 are in Supplementary Table 1. Linear regression results for drug effects versus replication timing are in Supplementary Table 2. Wilcoxon test results for the H3K27me2 and H3K27ac response to EPZ-6438 and EED-226 in S-phase fractions are in Supplementary Table 3.

## Supplemental Text and Figures

**Supplementary Figure 1.**
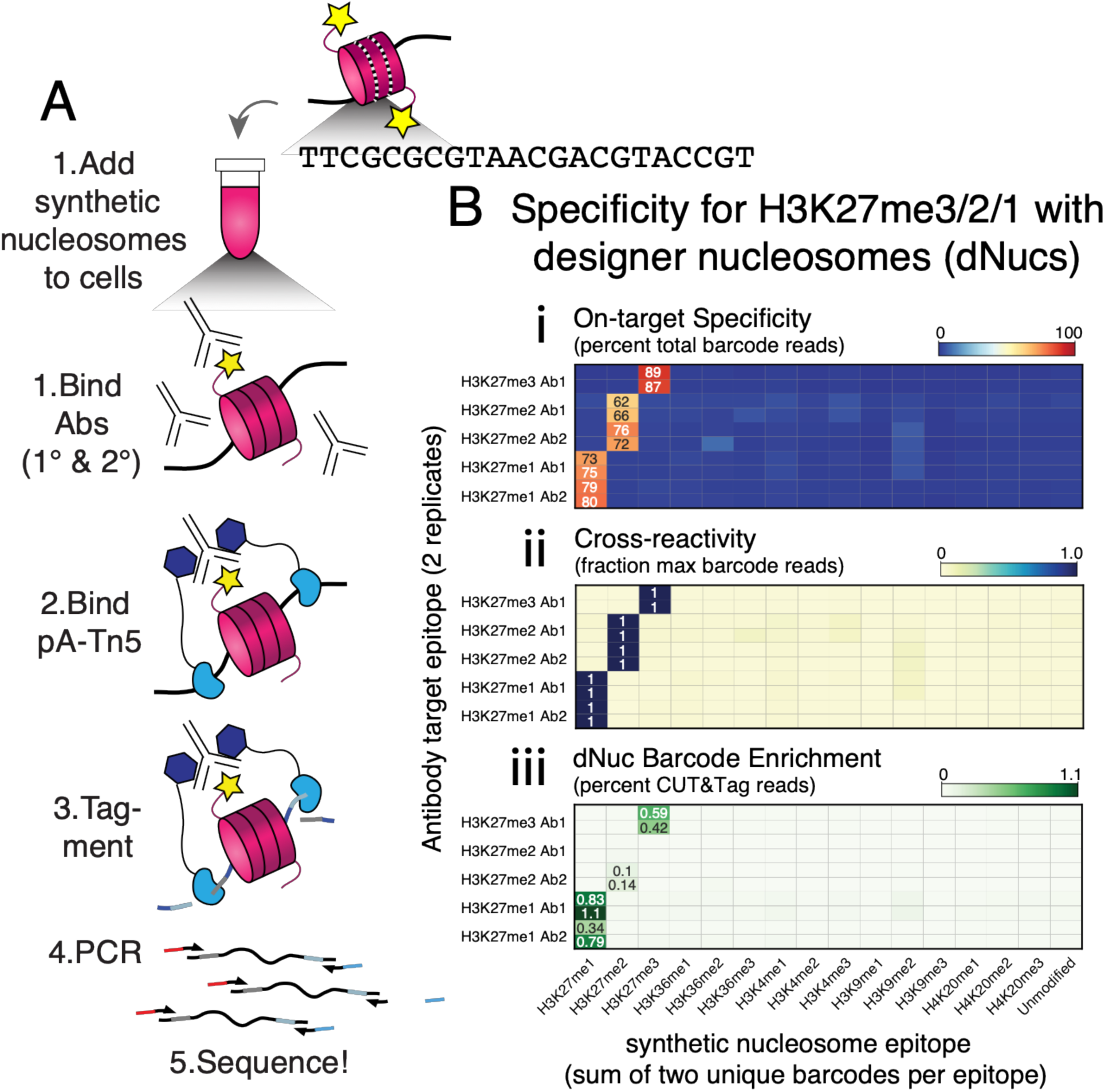
Confirmation of H3K27 methylation profiling with designer barcoded nucleosomes: (A) Cleavage Under Targets & Tagmentation (CUT&Tag) chromatin profiling paradigm with a synthetic nucleosome spike-in panel (EpiCypher, Cat# 19-1002). The synthetic nucleosomes are barcoded according to their lysine methylation status using a non-endogenous sequence that can quantified prior to reference genome alignment. The star denotes one of many possible lysine methylations also present in cells input to CUT&Tag. (B i) Heatmap of barcodes collated from FASTQ files of CUT&Tag libraries. Y-axis are individual libraries generated with a commercial antibody specific for H3K27me3, H3K27me2 (2 antibodies tested), or H3K27m1 (2 antibodies tested). Profiling was performed in duplicate. Values are the percent of total barcodes in the spike-in panel. (ii & iii) Same as (i) but values are (ii) fraction of the maximum barcode detected and (iii) percent total reads in the FASTQ file.

**Supplementary Figure 2.**
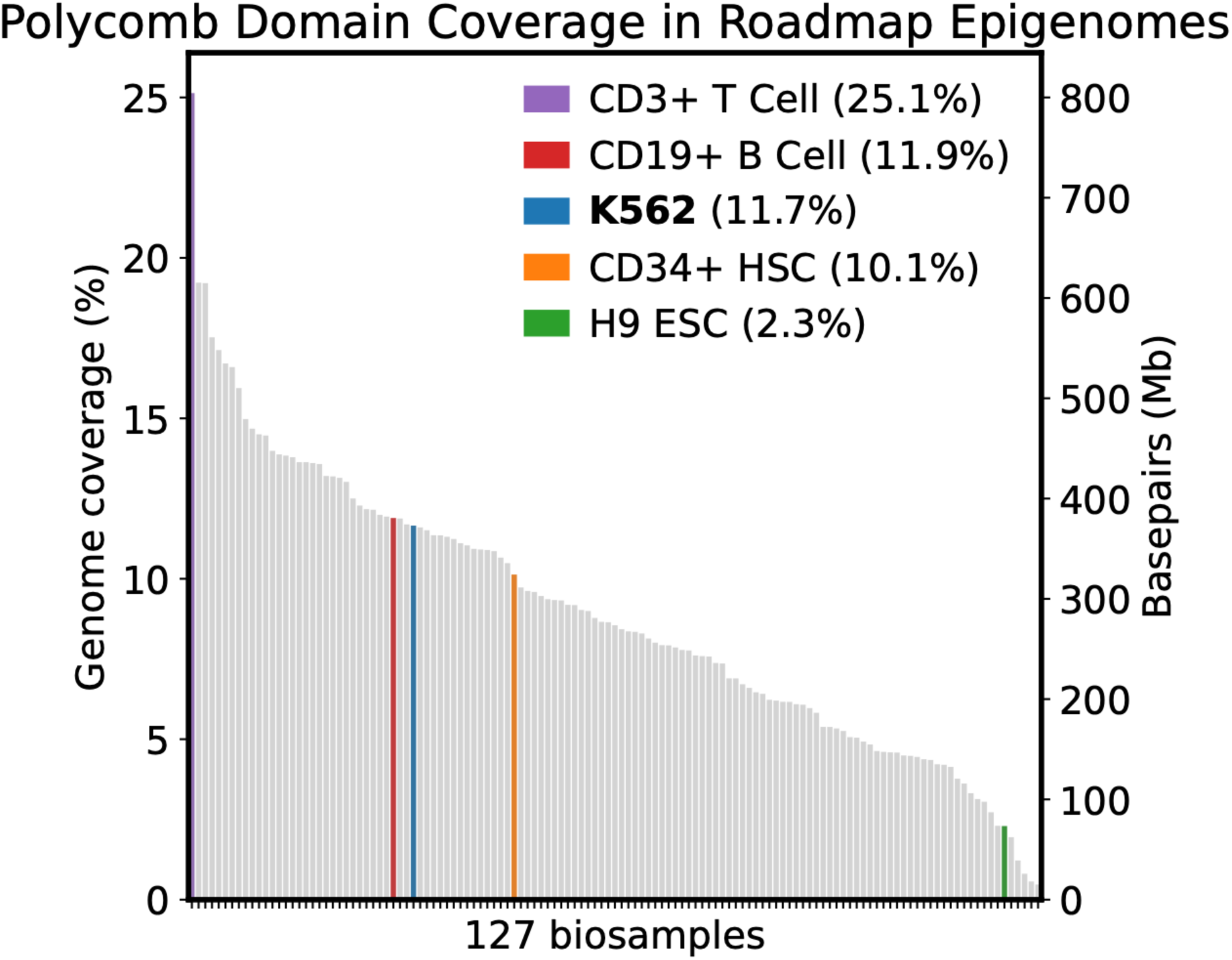
Polycomb domains defined by the Roadmap Epigenomes Consortium in 127 biosamples: Polycomb domains defined by the Roadmap Epigenomes Consortium^21^ in 127 biosamples ranked by genome coverage (left) and base pair coverage (right). K562 and four representative biosamples are highlighted with their percent genomic coverage. Polycomb domains occupy 11.7% of the genome in K562, which is near the median of the 127 Roadmap epigenomes.

**Supplementary Figure 3.**
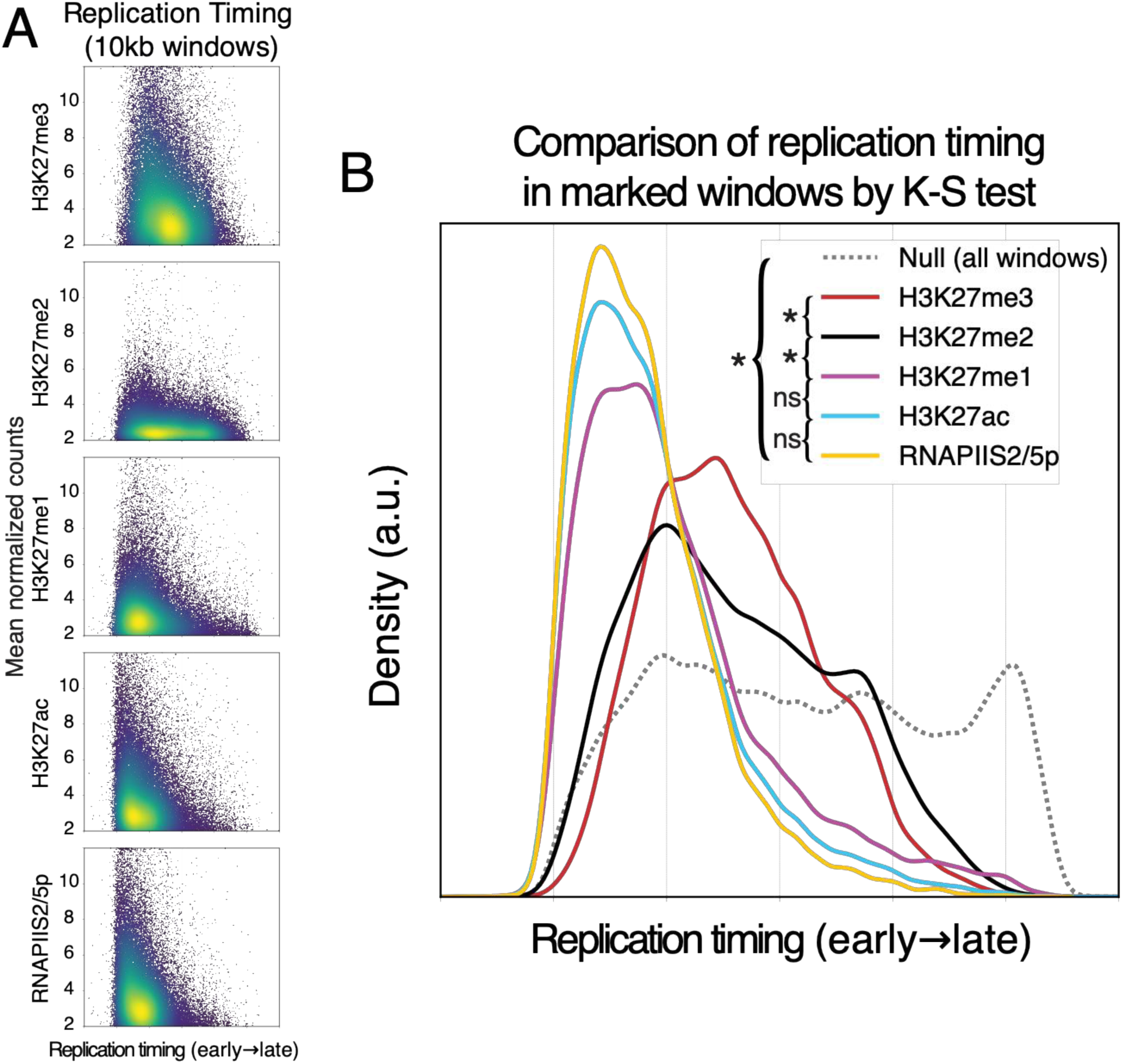
Replication timing of 10-kilobase (kb) windows with H3K27 modifications: (A) Mean normalized counts for H3K27me3, H3K27me2, H3K27me1, H3K27ac, and RNAPIIS2/5p (y-axis) in 10-kb windows versus mean replication timing (early-to-late) colored by 2d density (blue is low, yellow is high). A minimum threshold of 2 mean RPKM is applied to select for marked windows. (B) Dsitribution of replication timing in marked 10-kb windows compared across marks via the Kolmogorov-Smirnov test. The null distribution is replication timing in all 10kb windows. Comparisons are pairwise between marks and subject to Bonferroni correction. Comparisons between each mark and the null distribution were also performed (all significant). * = p < 0.01.

**Supplementary Figure 4.**
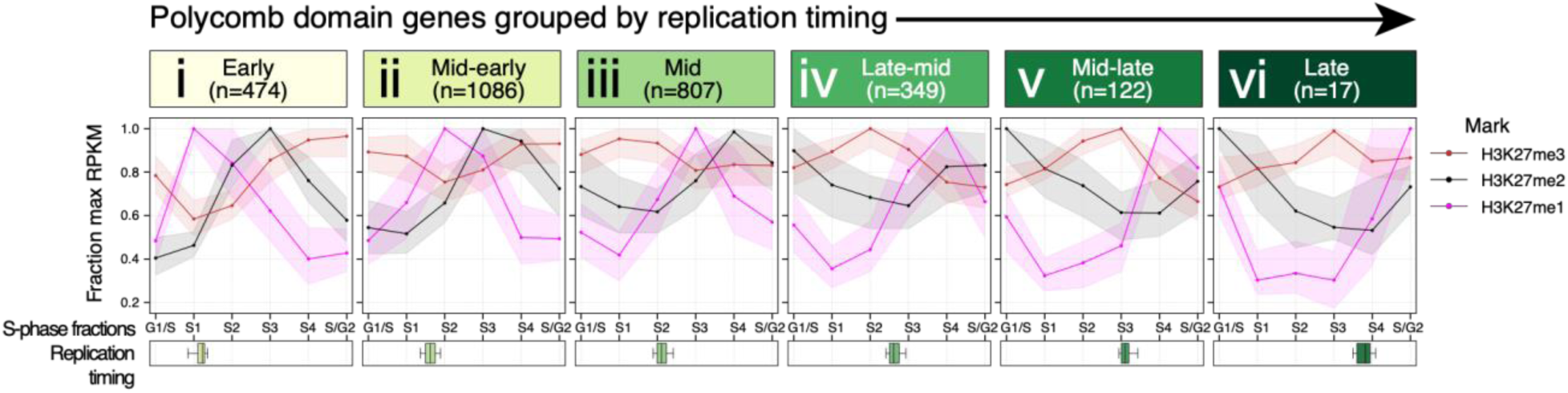
Trends of H3K27 methylation at Polycomb domain genes grouped by replication timing: Trends are median with interquartile range per S-phase fraction for H3K27me1 (magenta), H3K27me2 (black), and H3K27me3 (red) in Polycomb domain genes grouped by replication timing from left to right (i-vi). Replication timing is shown per group as one boxplot where each fraction is evenly spaced such that G1/S = 0, S1 = 0.2, S2 = 0.4, S3 = 0.6, S4 = 0.8, and S/G2 = 1.

**Supplementary Figure 5.**
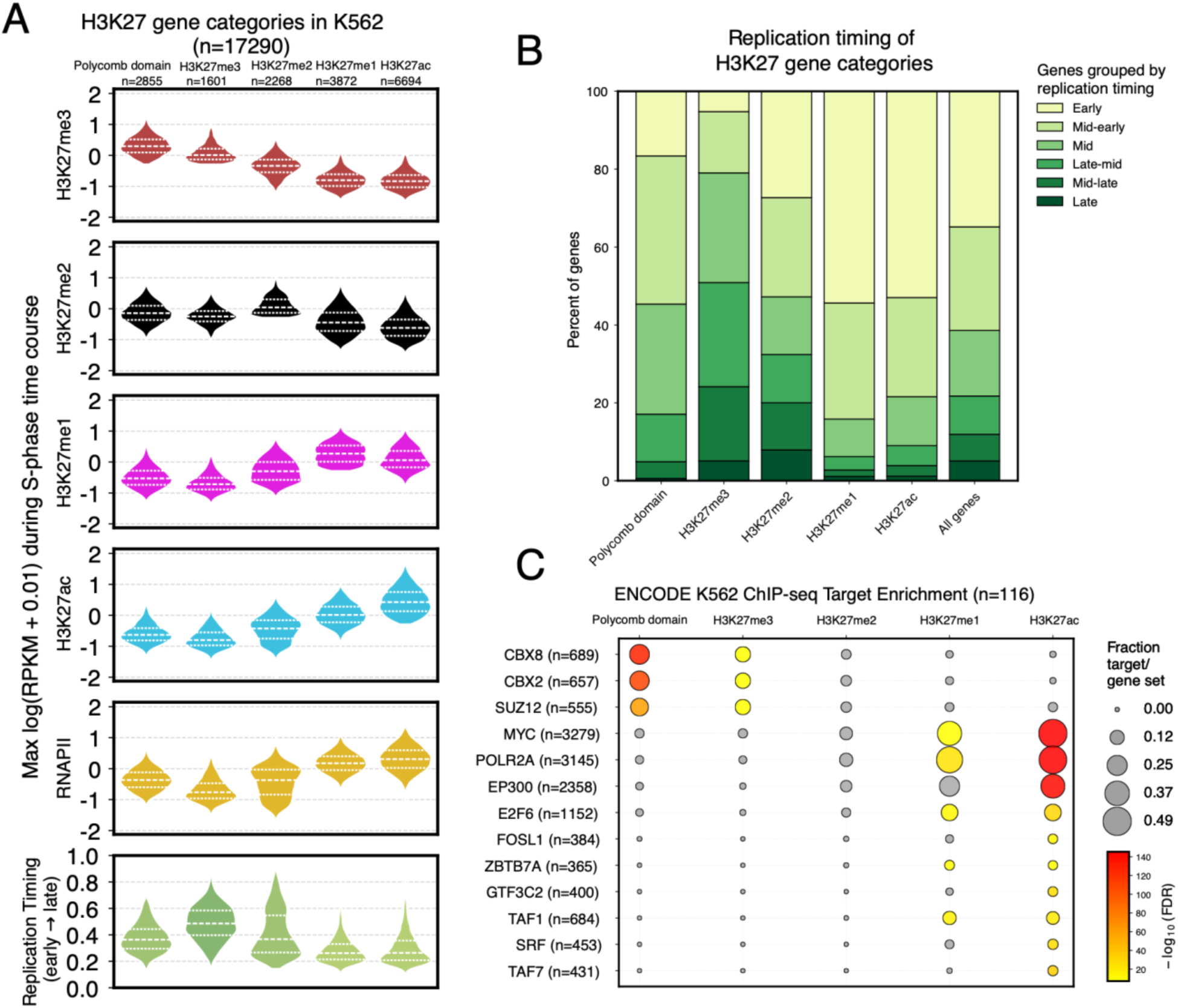
Assignment of H3K27 gene categories used in. **Figure 2**: (A) Genes in Polycomb domain (n = 2855) were assigned based on at least 50% base pair overlap with regions defined by the Roadmaps Epigenome Consortium^21^ using chromHMM^22^ in K562 cells. The remaining genes were then categorized based on the highest maximum H3K27 modification (RPKM) during the time course above a minimum of 0.5 RPKM. Signal is shown as log10(RPKM + 0.01) per gene, and replication timing is show from 0-1 (early-to-late) per category. (B) Replication timing groups per H3K27 category. (C) Overlap enrichment per H3K27 category versus ENCODE_TF_ChIP-seq_2015 target genes^59^ in K562. Top 3 gene sets (y-axis) are shown per H3K27 category.

**Supplementary Figure 6.**
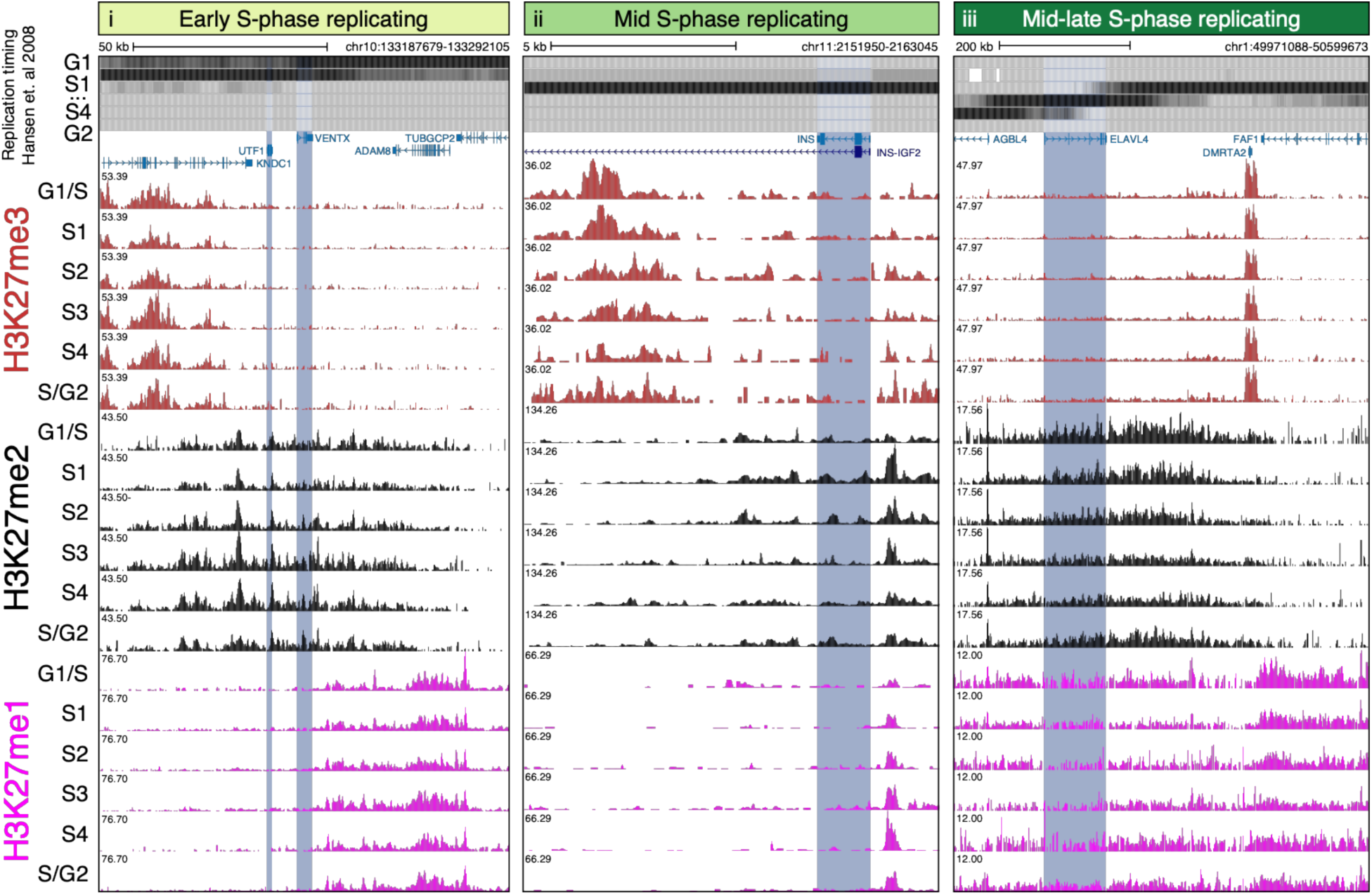
Examples of H3K27me2-enriched genes: (i) UCSC browser snapshot of coverage profiles for H3K27me3, H3K27me2, and H3K27me1 in S-phase fractions around the VENTX and UTF1 genes, highlighted in blue. Each mark is group auto-scaled. Replication timing in K562 cells^27^ is show in S-phase fractions at the top. Gene annotation is Gencode v49 for hg38. (ii-ii) same as (i) but (ii) at the INS gene and (iii) ELVAL4 gene.

**Supplementary Figure 7.**
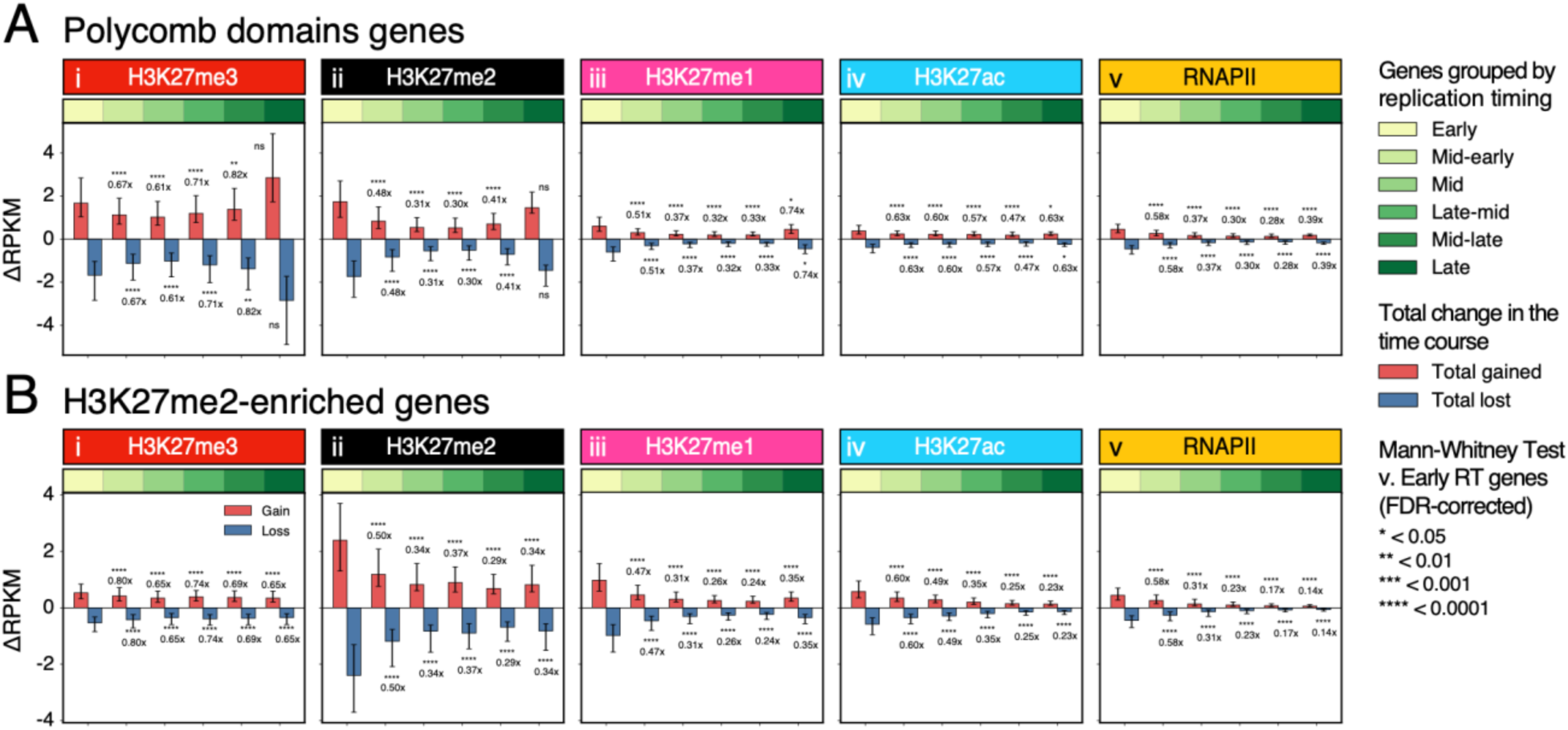
High change of H3K27 modification and active RNAPII at earl replicating genes in and outside Polycomb domains: (A) total change shown as delta-RPKM for (i) H3K27me3, (ii) H3K27me2, (iii) H3K27me1, (iv) H3K27ac, and (v) active RNAPII during the time course at genes in Polycomb domains grouped by replication timing (x-axis). Early replicating genes are compared to the other groups using the Mann-Whitney test and fold change is shown between medians. Error bars are interquartile range. (B) same as (A) but for H3K27me2-enriched genes.

**Supplementary Figure 8.**
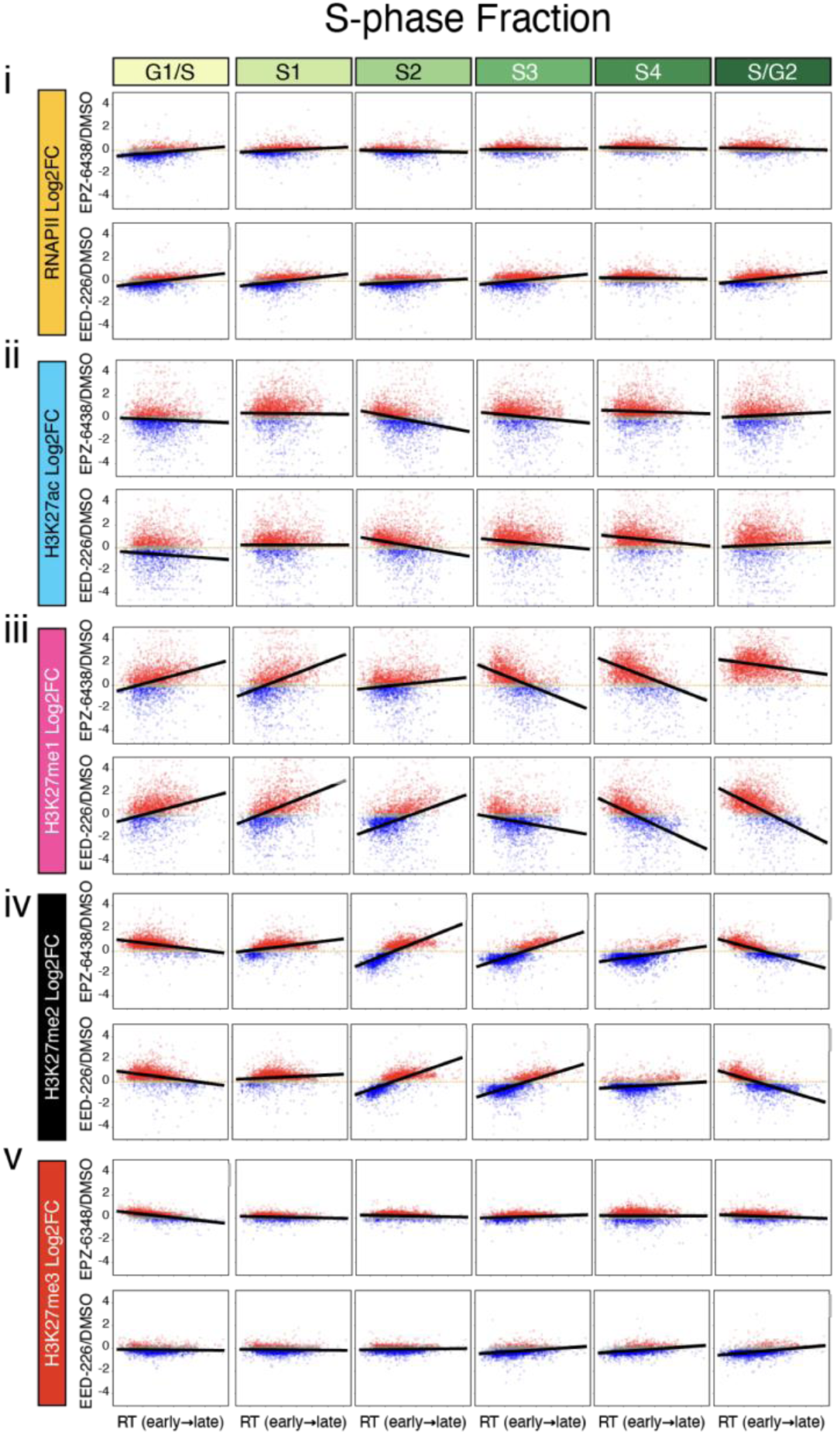
Linear regression of active RNAPII, H3K27ac, and H3K27 methylation in response to PRC2 inhibition versus replication timing at genes in Polycomb domains: (i) Log2 fold-change (y-axis) for EPZ-6438 (TAZ, top) and EED-226 (EED, bottom) versus DMSO is shown for RNAPII versus RT (x-axis, early-to-late) at genes in Polycomb domains (n=2861) per S-phase fraction (columns, left to right). Genes above an absolute of log2 fold-change of 0.2 are colored red for up in treatment or blue for down in treatment, or grey for Log2 fold-change < 0.2. The best fit line is shown per S-phase fraction. (ii-v) Same as (i) but for H3K27ac, H3K27me1, H3K27me2, and H3K27me3.

**Supplementary Figure 9.**
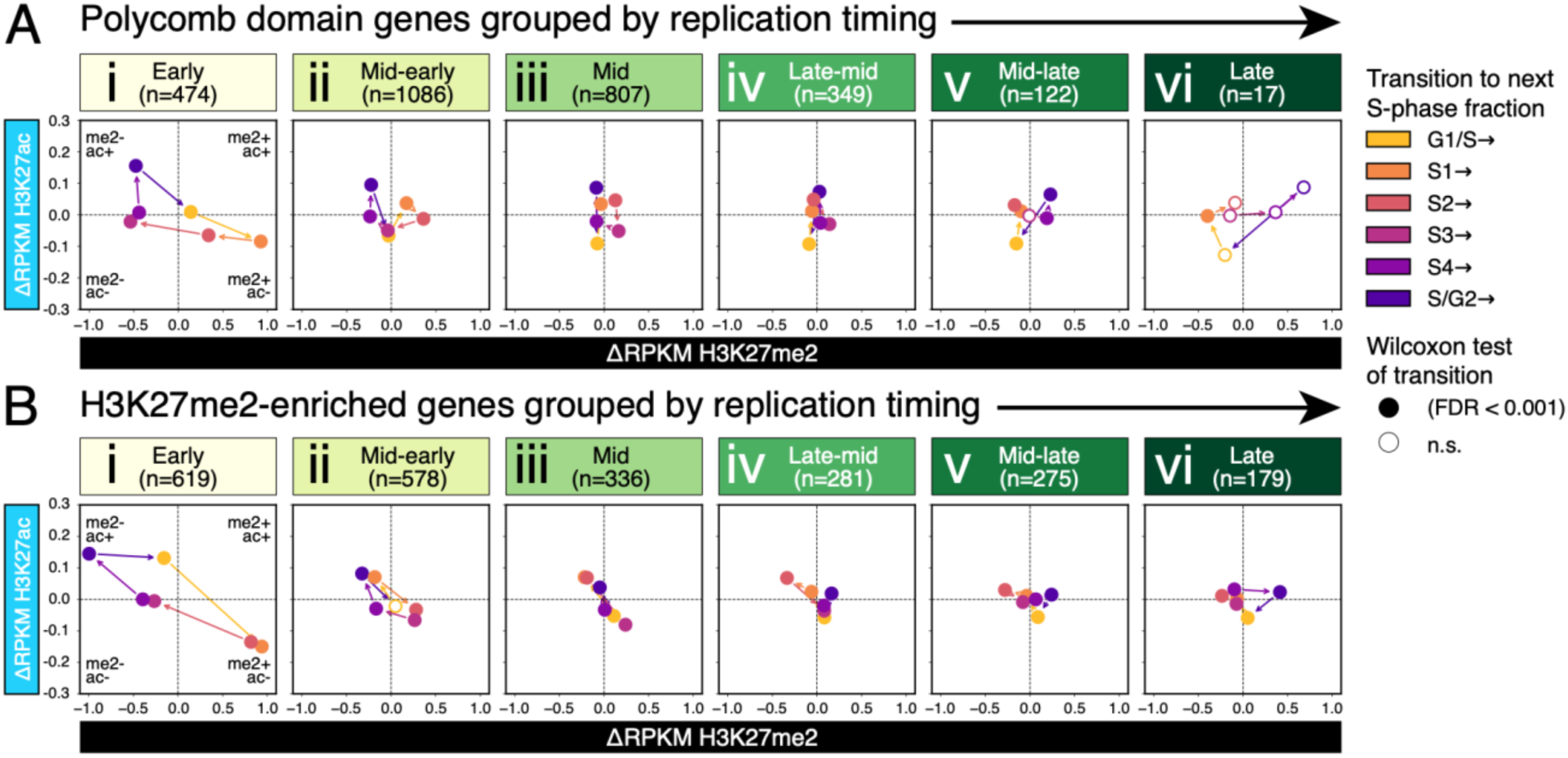
Changes in H3K27 acetylation versus di-methylation at genes in and outside Polycomb domains without drug treatment: (A) Magnitude of change shown as delta-RPKM between S-phase fractions for active H3K27ac versus H3K27me2 at genes in Polycomb domains grouped by replication timing. Filled dots are medians with a significant change in either mark. Arrows show the direction of the cell cycle time course between S-phase fractions. (B) Same as (A) but for H3K27me2-enriched genes.

**Supplementary Figure 10.**
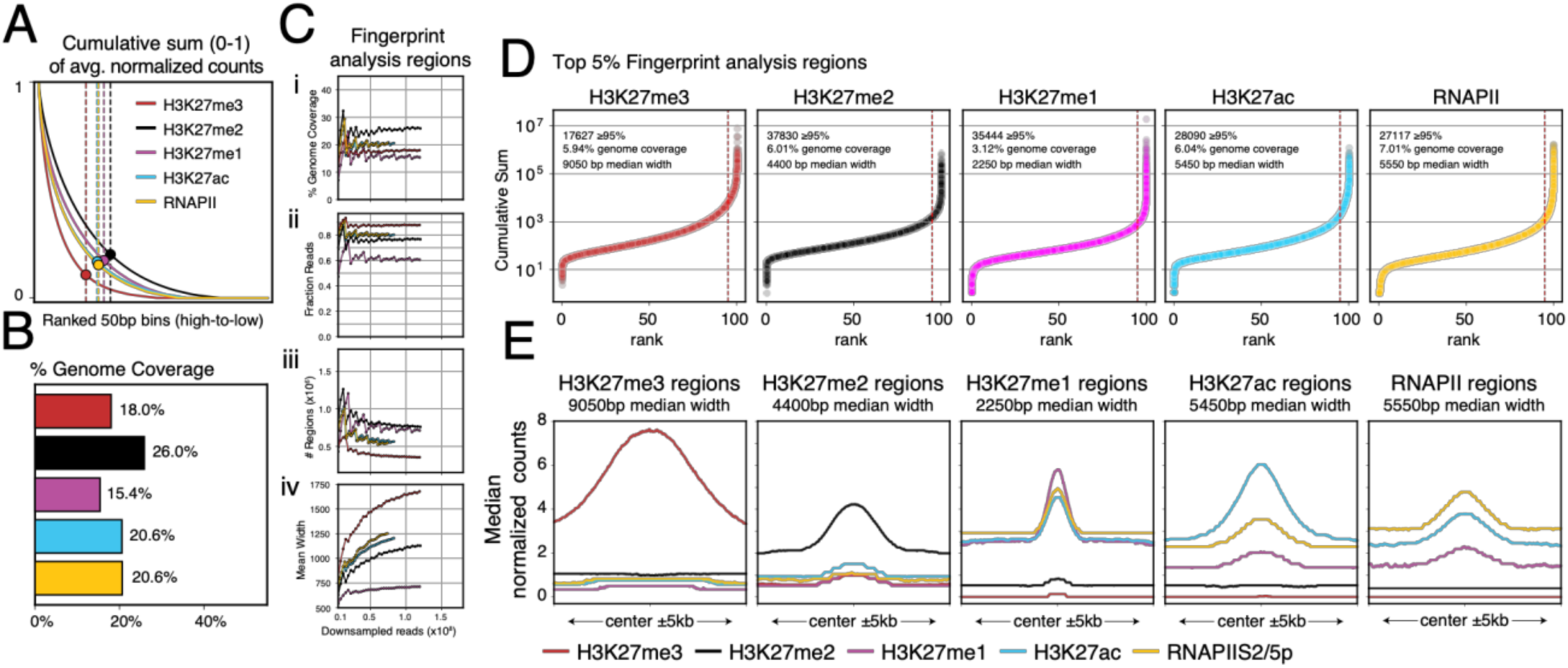
Definition of H3K27-marked regions for TRIP analysis in Figure 5: Fingerprint analysis^60^ of H3K27 modifications and active RNAPII steady-state profiles performed in 50-basepair (bp) intervals. Adjacent bins above a slope threshold of 1 were merged, and intervals <300bp were culled. This approach yielded consistent results down to ∼10 million reads per mark. (A) Cumulative sum in ranked 50bp bins. Slope threshold of 1 is shown as a dotted line. (B) Percent genome covered by intervals > 300bp per mark. (C) Benchmarking of fingerprint analysis regions via down sampling the reads used to make the coverage file. Metrics (y-axis) versus down sampled reads (x-axis) are shown from top-to-bottom: (i) % genome coverage, (ii) fraction of reads in fingerprint regions, (iii) # of regions, and (iv) mean region width. (D) Regions ranked by the sum of normalized counts. Statistics are shown for the top 5% of intervals. (E) Median normalized coverage of H3K27me3, H3K27me2, H3K27me1, H3K27ac, and RNAPIIS2/5p centered over the top 5% of regions per mark. Regions that did not overlap those of other marks were used for TRIP analysis, except for regions common to H3K27me1 and H3K27ac.

**Supplementary Figure 11.**
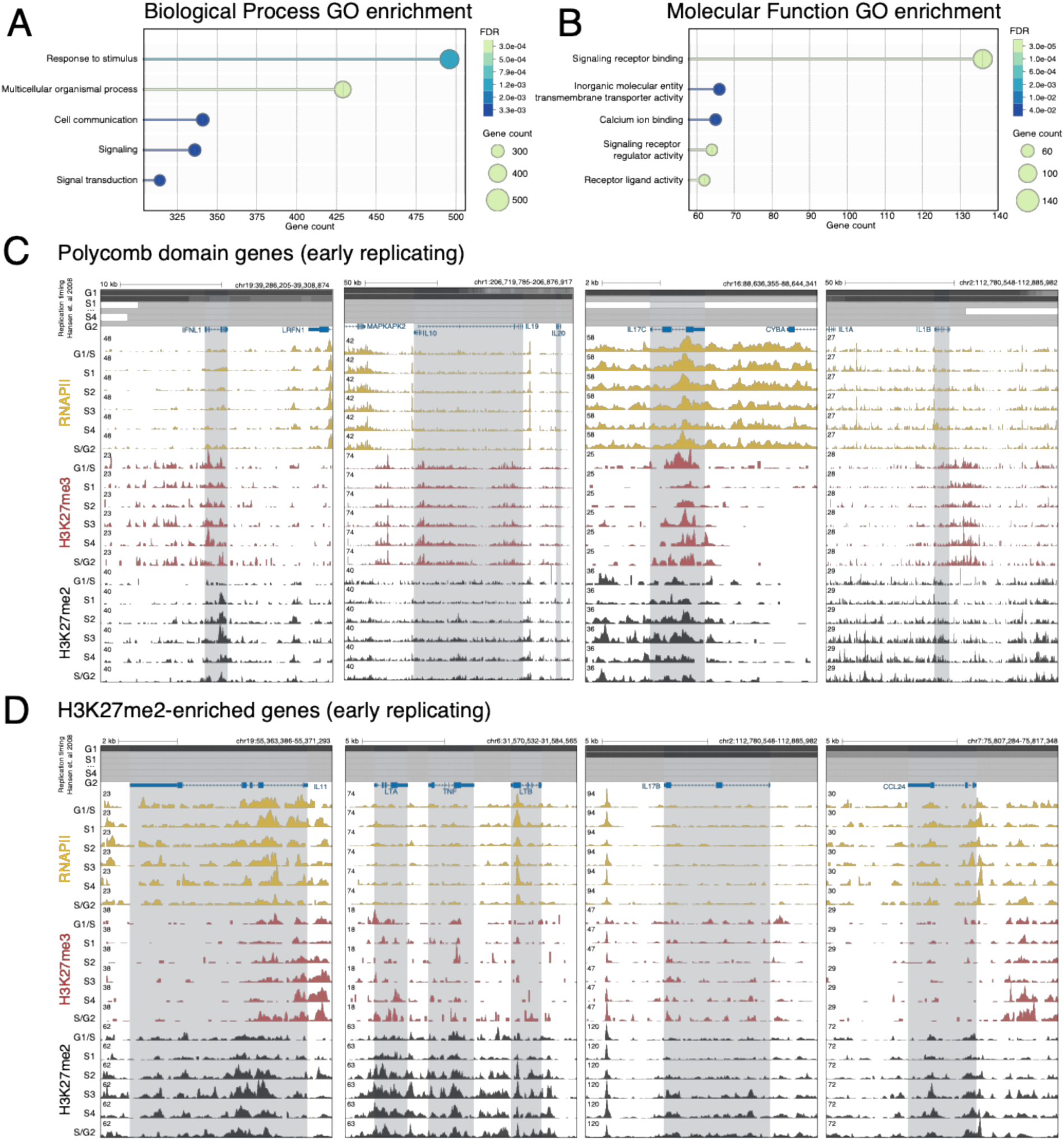
Description and examples of early replicating genes in and outside Polycomb domains: (A-B) Gene ontology (GO) enrichment analysis of early replicating genes in and outside Polycomb domains for (A) Biological process and (B) Molecular function generated using the String database^61^. Categories were ranked by number of overlapping genes. (C-D) Examples of (C) Polycomb domain genes and (D) H3K27me2-enriched genes in the ‘Response to stimulus’ Biological process GO category that encode cytokine signaling molecules. UCSC browser snapshots show of coverage profiles for Active RNAPII (yellow), H3K27me3 (red), and H3K27me2 (black) in S-phase fractions around the gene loci, highlighted in blue. Each mark is group auto-scaled. Replication timing in K562 cells^27^ is show in S-phase fractions at the top. Gene annotation is Gencode v49 for hg38. Genes in (C) are *IFNL1, IL19, IL10, IL17C, IL20, and IL10*. Genes in (D) are *ILII, LTA, TNF, LTB, IL17B, and CCL24*.

